# Transcription factor network analysis identifies REST/NRSF as an intrinsic regulator of CNS regeneration

**DOI:** 10.1101/2020.12.13.413104

**Authors:** Yuyan Cheng, Yuqin Yin, Alice Zhang, Alexander M. Bernstein, Riki Kawaguchi, Kun Gao, Kyra Potter, Hui-Ya Gilbert, Yan Ao, Jing Ou, Catherine J. Fricano-Kugler, Jeffrey L. Goldberg, Clifford J. Woolf, Michael V. Sofroniew, Larry I. Benowitz, Daniel H. Geschwind

## Abstract

The inability of neurons to regenerate long axons within the CNS is a major impediment to improving outcome after spinal cord injury, stroke, and other CNS insults. Recent advances have uncovered an intrinsic program that involves coordinate regulation by multiple transcription factors that can be manipulated to enhance growth in the peripheral nervous system. Here, we used a system-genomics approach to characterize regulatory relationships of regeneration-associated transcription factors, identifying RE1-Silencing Transcription Factor (REST; Neuron-Restrictive Silencer Factor, NRSF) as a predicted upstream suppressor of a pro-regenerative gene program associated with axon regeneration in the CNS. We validate our predictions using multiple paradigms, showing that mature mice bearing cell type-specific deletions of REST or expressing dominant-negative mutant REST showed improved regeneration of the corticospinal tract and optic nerve, accompanied by upregulation of regeneration-associated genes in cortical motor neurons and retinal ganglion cells, respectively. These analyses identify a novel role for REST as an upstream suppressor of the intrinsic regenerative program in the CNS and demonstrate the power of a systems biology approach involving integrative genomics and bio-informatics to predict key regulators of CNS repair.

## INTRODUCTION

Injured axons in the adult mammalian central nervous system (CNS) generally cannot regenerate over long distances, limiting functional recovery from CNS injury (*1*). Potential mechanisms underlying regenerative failure in the mature CNS include a lack of an intrinsic ability to activate genes and pathways required for axon regrowth after injury (*2, 3*); the presence of extrinsic growth-repulsive factors associated with certain extracellular matrix molecules, myelin debris, or fibrotic tissue (*4–6*); and limited availability of appropriate growth factors (*1, 7, 8*). Strategies to neutralize or attenuate key cell-extrinsic inhibitors of axon growth have limited effects on regeneration (*9, 10*), though their impact is strongly enhanced by co-activating neurons’ intrinsic growth state (*11–13*). Deleting PTEN, a cell-intrinsic suppressor of axon growth, induces appreciable axon regeneration, and when combined with either CNTF plus SOCS3 deletion, or with inflammation-associated factors plus cAMP, enables a percentage of retinal ganglion cells to regrow axons the full length of the optic nerve (*14–17*). Nonetheless, more work is needed to identify key regulators of axon regeneration in the CNS, including transcription factors that act as master switches of the regenerative program.

Unlike their CNS counterparts, peripheral sensory and motor neurons spontaneously display potent growth in response to peripheral axonal injury, which is accompanied by activation of key regeneration-associated genes (RAGs) (*18, 19*) that we recently found to act as a coordinated network to promote growth (*20*). Expression of this RAG network is predicted to be regulated by a core group of TFs during peripheral nerve regeneration (*20*). This prediction is supported by the findings that manipulating individual TFs at the core of this network, such as STAT3 (*21*), KLF family members (*22, 23*), and Sox11 (*24, 25*) result in varying amounts of CNS axon growth. The effects of TFs on their target pathways are dynamic, combinatorial, and form tiered regulatory networks, requiring tight control in timing, dosage, and the context of each TF involved (*26–30*). The complexity of recapitulating coordinated TF regulatory events may limit the effectiveness of single gain- or loss-of-function experiments to determine contributions of individual TFs within a complex network (*31*). Alternatively, illuminating the hierarchical transcriptional network architecture from gene expression datasets provides an efficient means to identify key upstream regulators of various biological processes, for example pluripotency (*32*). Predominant models of TF networks rely on a 3-level pyramid-like structure, with a small number of TFs at the top-level that function as ‘master’ regulators, driving expression of most of the other mid- and bottom level TFs that directly or indirectly regulate the expression of their target genes (*30, 33–35*).

Here, we integrated multiple existing and newly-generated datasets to characterize hierarchical TF interactions so as to identify potential upstream regulators associated with the intrinsic axon regeneration state (Figure 1A). By comparing gene expression in non-permissive states, such as the injured CNS, to the permissive PNS or to the CNS that has been subjected to strong pro-regenerative treatments, we hypothesized that we could identify key upstream TFs driving intrinsic regeneration programs. We began with a mutual information-based network analysis approach to characterize the transcriptional regulatory network formed by regeneration-associated TFs (*20*) in multiple independent data sets. We identified a core three-level subnetwork of five interconnected TFs, consisting of Jun, STAT3, Sox11, SMAD1, and ATF3, which is strikingly preserved across multiple PNS injury models and at different timescales (*36–39*). Remarkably, we observe a similar multi-layer, highly inter-connected TF structure in CNS neurons following genetic and pharmacological treatments that enhance regeneration. In contrast, in the non-regenerating CNS at baseline (*36, 40, 41*), this regeneration-associated subnetwork and its 3-tier hierarchical structure are dismantled, and candidate TFs adopt a less interconnected and less hierarchical structure.

**Figure 1.**
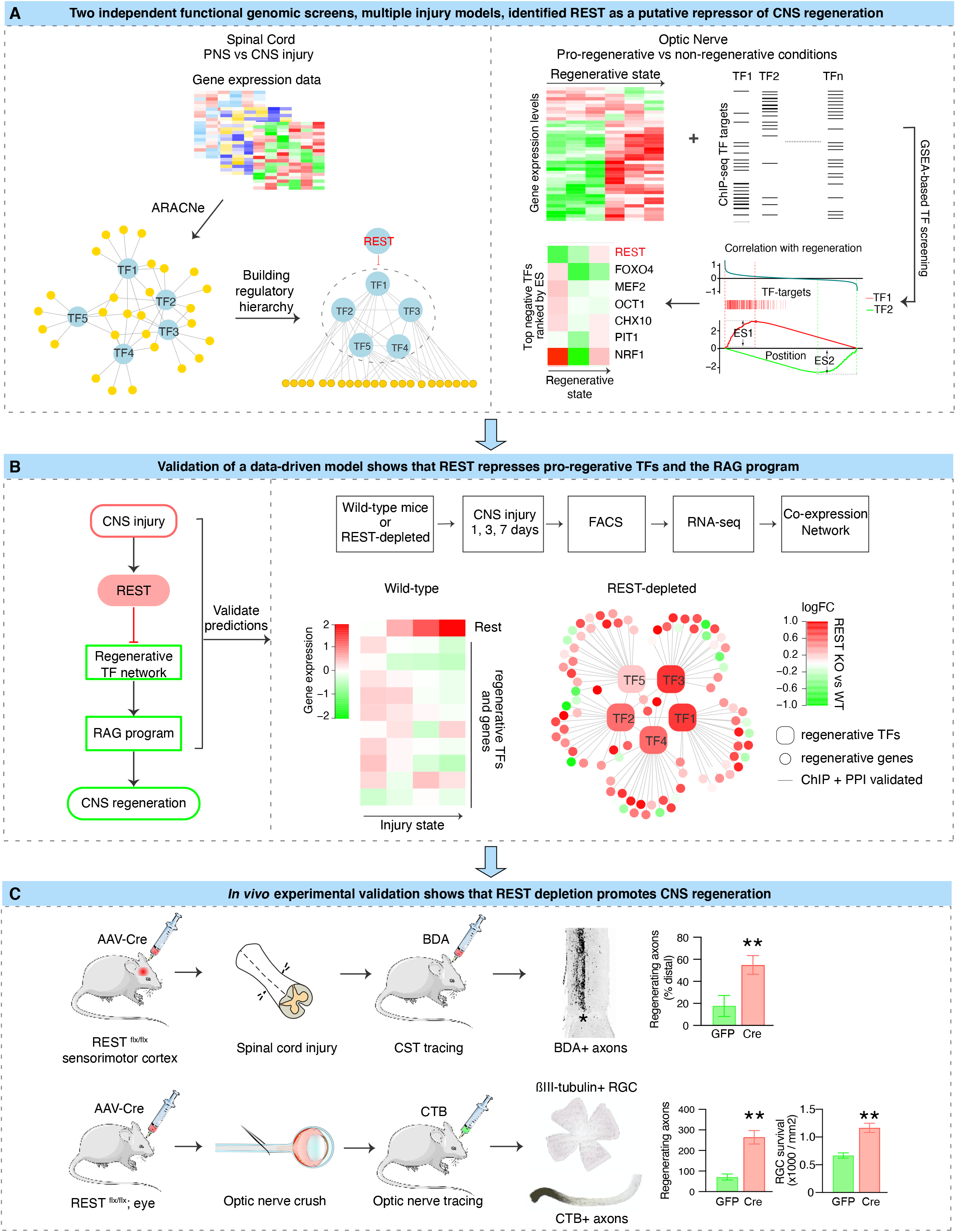
Schematic diagram summarizing the overall experimental flow integrating iterative bio-informatics and experimental validation. Multiple independent functional genomics analyses of distinct injury models were analyzed to computationally identify upstream TFs associated with CNS regeneration. In the first set of analysis (A, left), we performed a mutual information-based network analysis using ARACNe to characterize the transcriptional regulatory network formed by regeneration-associated TFs in multiple independent data sets from spinal cord and peripheral nerve injury. The hierarchical structure of the TF regulatory network was further characterized, so as to identify potential upstream regulators. This step-wise analysis predicted REST, a transcriptional repressor, as an upstream negative regulator inhibiting the core pro-regenerative TFs to drive the expression of regeneration-associated genes (RAGs). In parallel (A, right), we performed an additional unbiased genome-wide screen in another CNS tissue, optic nerve, under pro-growth and native conditions to identify TF regulators of regeneration. Among the ∼1000 TF-target gene sets unbiasedly tested via Gene Set Enrichment Analysis, REST was ranked as the top negative regulator of the RGC regeneration state-associated gene set. Multiple independent bio-informatic analyses of external data sets confirmed and converged on our model (B), by which REST is activated by CNS injury and acts as a potential upstream negative regulator of the core regenerative TFs. To test this, we performed gene expression analysis in the injured CNS with REST and after REST depletion, showing REST increases following CNS injury, while the core pro-regenerative TFs and genes remain suppressed. Depleting REST activates a core molecular program driven by a tightly controlled TF network similar to the one activated during regeneration. These results predicted that REST depletion would improve regeneration, which we directly tested in two different, well-established models of regeneration *in vivo* (C), confirming REST’s functional effect as a suppressor of regeneration. In the case of optic nerve injury, REST depletion or inhibition enhanced both RGC regeneration and survival. These analyses identify a novel role for REST as an upstream suppressor of the intrinsic regenerative program in the CNS and demonstrate the power of a systems biology approach involving integrative genomics and bio-informatics to predict key regulators of CNS repair.

Our analyses identified RE1-silencing transcription factor (REST; (*42, 43*)), a widely studied regulator of neural development and neural-specific gene expression (*42–45*) (*46, 47*), as playing a potentially important role in suppressing CNS regeneration (Figure 1). The bio-informatic analyses showed that REST, a repressive factor, is present at the apex of a degenerate TF network in the non-regenerating CNS, but absent in the PNS and in CNS neurons with enhanced regenerative potential, both in the optic nerve and spinal cord. Our findings suggested that REST acts as a potential upstream transcriptional repressor, limiting the interactions of the core regenerative TFs to drive the expression of RAGs and the intrinsic growth capacity of CNS neurons (Figure 1B). This hypothesis was supported by transcriptomic analysis of REST-depleted, CNS-injured neurons, which displayed enhanced expression of a regeneration-associated gene network, driven by several core TFs known to promote regeneration. To further validate our bio-informatic predictions, we investigated the effects of counteracting REST on regeneration in two different models of CNS injury *in vivo* – optic nerve crush and complete spinal cord injury (SCI) – via conditional depletion or functional inactivation of REST in retinal ganglion cells (RGCs) and corticospinal tract (CST) projection neurons (Figure 1C). In both cases, counteracting REST resulted in increased regeneration. These findings demonstrate how a multi-step systems biological analysis coupled with substantial *in vitro* and *in vivo* experimental validation provides a framework for discovery of drivers of CNS repair, and implicate REST as a novel regulator of CNS axon regeneration.

## RESULTS

### Bio-informatic analysis identifies REST as a potential upstream repressor of a regeneration-associated network

To determine which of the previously identified pro-regenerative TFs (*20*) are essential drivers of the neural intrinsic growth program, we characterized the regulatory network among these TFs to define their directional and hierarchical relationships. We employed a step-wise approach, summarized in Figure 2A. To infer directionality of each pair of TFs, we applied the Algorithm for Reconstruction of Accurate Cellular Networks (ARACNe), a mutual-information (MI) based algorithm for reverse-engineering transcriptional regulatory network from gene expression datasets (*48, 49*). ARACNe connects two genes only if there is an irreducible statistical dependency in their expression. These connections likely represent direct regulatory interactions mediated by a TF binding to its target genes, which could be TFs, and thus can be used to predict the TF network and their transcriptional targets (*48*) (Figure 2A; Methods). These predictions have been extensively validated by experimental analysis, such as chromatin immunoprecipitation sequencing (ChIP-Seq), a method to identify physical TF-target binding, or by examining expression changes of target genes led by gain- or loss- of function of the regulatory TFs (*50–54*).

**Figure 2.**
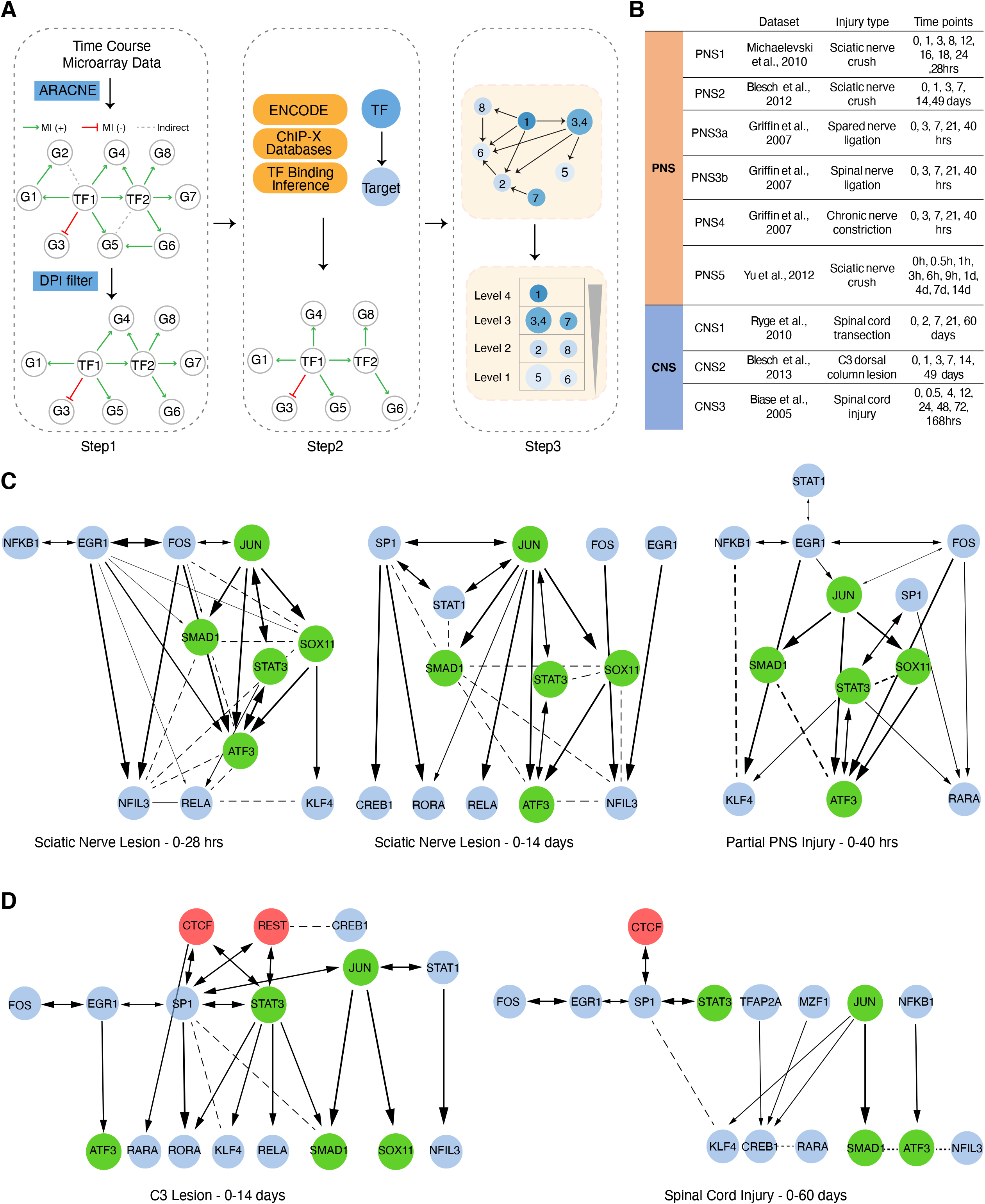
Characterizing regeneration-associated transcriptional regulatory network. **(A)** Schematic diagram illustrating step-wise approaches employed to infer hierarchical TF regulatory networks from **(B)** time-course microarray datasets. Step 1: First, ARACNe was applied to each dataset to find TF-target pairs that display correlated transcriptional responses by measuring mutual information (MI) of their mRNA expression profiles (Methods). The sign (+/-) of MI scores indicates the predicted mode of action based on the Pearson’s correlation between the TF and its targets. A positive MI suggests activation of this TF on its targets, while a negative MI score suggests repression. All non-significant associations were removed by permutation analysis. Second, ARACNe eliminates indirect interactions, such as two genes connected by intermediate steps, through applying a well-known property of MI called data-processing inequality (DPI). Step 2: To determine the direction of regulation between each TF interactions, ChIP-datasets from ENCODE and previously published ChIP-ChIP and ChIP-seq datasets were integrated to compile a list of all observed physical TF-target binding interactions. Step 3: To identify the hierarchical structure within directed TF networks, we used graph-theoretical algorithms to determine precise topological ordering of directed networks based on the number of connections that start from or end at each TF, indicating whether a TF is more regulating or more regulated. **(C-D)** Representative regulatory networks inferred from microarrays following peripheral nerve injury **(C)** and CNS injury **(D)**. Each node represents one of the 21 regeneration-promoting TFs if a connection exists. The thickness of each line indicates the MI between the TFs it connects. A directional arrow is drawn if there is direct physical evidence of the TF binding its target TF’s promoter.

In an ARACNe-constructed transcriptional regulatory network, a TF is either predicted to have a positive edge with its target genes (i.e. activator of expression; MI (+)) when their expression patterns are positively correlated, or negative edge (i.e. repressor of expression, MI (-)) if the TF displays opposite transcriptional changes from its targets (Figure 2A, step 1). We subsequently validated the initial bio-informatic predictions of edge directionality by compiling direct biochemical evidence of physical TF-target binding observed by multiple ChIP-Seq or ChIP-ChIP databases (*55, 56*), leading to a high-confidence, directed TF regulatory network supported by experimental evidence (Figure 2A, step 2). Lastly, the hierarchical structure of the directed TF network was defined by a graph-theoretical algorithm (*33*), which constructs the precise topological ordering of members in any directed network (*35*) (Figure 2A, step 3).

Because TF binding is a dynamic process that may change over time, we analyzed 9 high-density time-series gene expression profiles upon injury to build the networks, leveraging the chronological order of regulatory events. By applying our step-wise pipeline to 6 peripheral nerve and 3 spinal cord injury datasets (Figure 2B; Methods), we sought to identify reproducible differences in transcriptional regulatory networks between regenerating PNS and non-regenerating CNS neurons following injury. We found that the candidate TFs regulate each other within complex, multi-layered networks, similar to TF network models defined by ENCODE (Figure 2C; (*30, 34*)). Across multiple datasets in multiple PNS injury models collected in different laboratories at different timescales, we observed a remarkable preservation of a defined five TF subnetwork, consisting of JUN, STAT3, SOX11, SMAD1, and ATF3 (Figure 2C), all of which are required for peripheral nerve regeneration (*20, 57–66*). In striking contrast, this subnetwork is dismantled and adopts a simpler, bi-layered, less inter-connected, and less hierarchical structure in the case of CNS injury (Figure 2D).

The expression levels of all 5 core TFs (*Atf3, Jun, Sox11, Stat3, Smad1*) were consistently increased among multiple PNS injury datasets (Figure S1A). Their increases occur as early as 0.5-3 hours after PNS injury (Figure S1A, PNS1, 3, and 5), and are maintained for as long as ∼40 days (Figure S1A, PNS2, and 4). In contrast, in the CNS, these key TFs were either not induced by CNS injury (Figure S1A, CNS1 and 2), or were transiently up-regulated but quickly repressed at later stage (Figure S1A, CNS3). In addition, *Atf3*, *Jun*, *Sox11*, and *Smad1* bear the most correlated regulatory relationships with others across multiple PNS injury datasets (Figure S1B PNS vs PNS). By contrast, there is little correlation in the regulatory interactions of the core TFs between PNS and CNS injury datasets (Figure S1B, CNS vs PNS). This finding is in general agreement with previous work (*20*), in which the peripherally activated RAG program predicted to be targeted by these TFs is highly preserved in PNS injury datasets, but not in the CNS. Our results indicate that a highly reproducible TF network potentially driving the expression of a RAG program is induced during peripheral nerve regeneration, but is significantly attenuated in the injured CNS.

Remarkably, we observed that two TFs, REST and CTCF, exhibit significant interactions with top-tier TFs in the CNS network, but not in the PNS network, and are predicted to inhibit other top-tier TFs. *Rest* mRNA levels did not change after PNS injury (Figure S1A, PNS1-5), but were increased by CNS injury when other key regenerative TFs began to be repressed (Figure S1A, CNS1-3). We did not observe changes of *Ctcf* expression levels following PNS or CNS injury. We therefore hypothesized that REST, which appears at the apex of a dismantled, less inter-connected TF network, was a potential upstream transcriptional repressor of the core TF network specifically in the non-regenerating CNS, thus limiting interactions between the core TFs to drive the expression of regeneration-associated genes and to activate the intrinsic growth state of CNS neurons.

### REST deletion in CNS-injured neurons increases expression of growth-related genes and pathways

We hypothesize that if REST were indeed an upstream repressor, as predicted by our bio-informatic model, its depletion in CNS neurons should release the transcriptional brake of pro-regenerative TFs and genes, subsequently increasing their expression. To test this hypothesis, we performed RNA-seq on REST-depleted sensorimotor cortical neurons that give rise to the corticospinal tract (CST) axons that course through the spinal cord. The CST is essential to control voluntary motor movements, and the failure of CST axons to regenerate is a major impediment to improving outcome after spinal cord injuries (*67*). To induce neuron-specific REST depletion, we injected adeno-associated virus expressing Cre recombinase or GFP as a control under a synapsin promoter (AAV-Syn-Cre and AAV-Syn-GFP) in the sensorimotor cortex of mice with homozygous conditional REST alleles and a TdTomato reporter (REST^flx/flx^; STOP^flx/flx^ TdTomato mice; Methods). REST knock-out (cKO) was confirmed by tdTomato expression in the cortical area of REST^flx/flx^; STOP^flx/flx^TdTomato mice injected with AAV-Syn-Cre. No TdTomato was observed in control mice receiving AAV-Syn-GFP. We then performed anatomically complete spinal cord crush at thoracic level 10 (T10) to avoid the spontaneous axon regeneration due to circuit reorganization that can occur after incomplete injury (*68*). Following sham or T10 SCI, neurons expressing GFP-(wild-type) or tdTomato-(REST cKO) were FACS-sorted at multiple time points post injury for RNA sequencing (Figure 3A; Methods). We then analyzed transcriptional differences in response to SCI and REST depletion at both the individual gene expression level and co-expression network level. Integrating network-level analysis complements analysis of differential expression by reducing the dimensionality of a large transcriptomic dataset and helps to find clusters of genes sharing expression patterns and biological functions (*69*).

**Figure 3.**
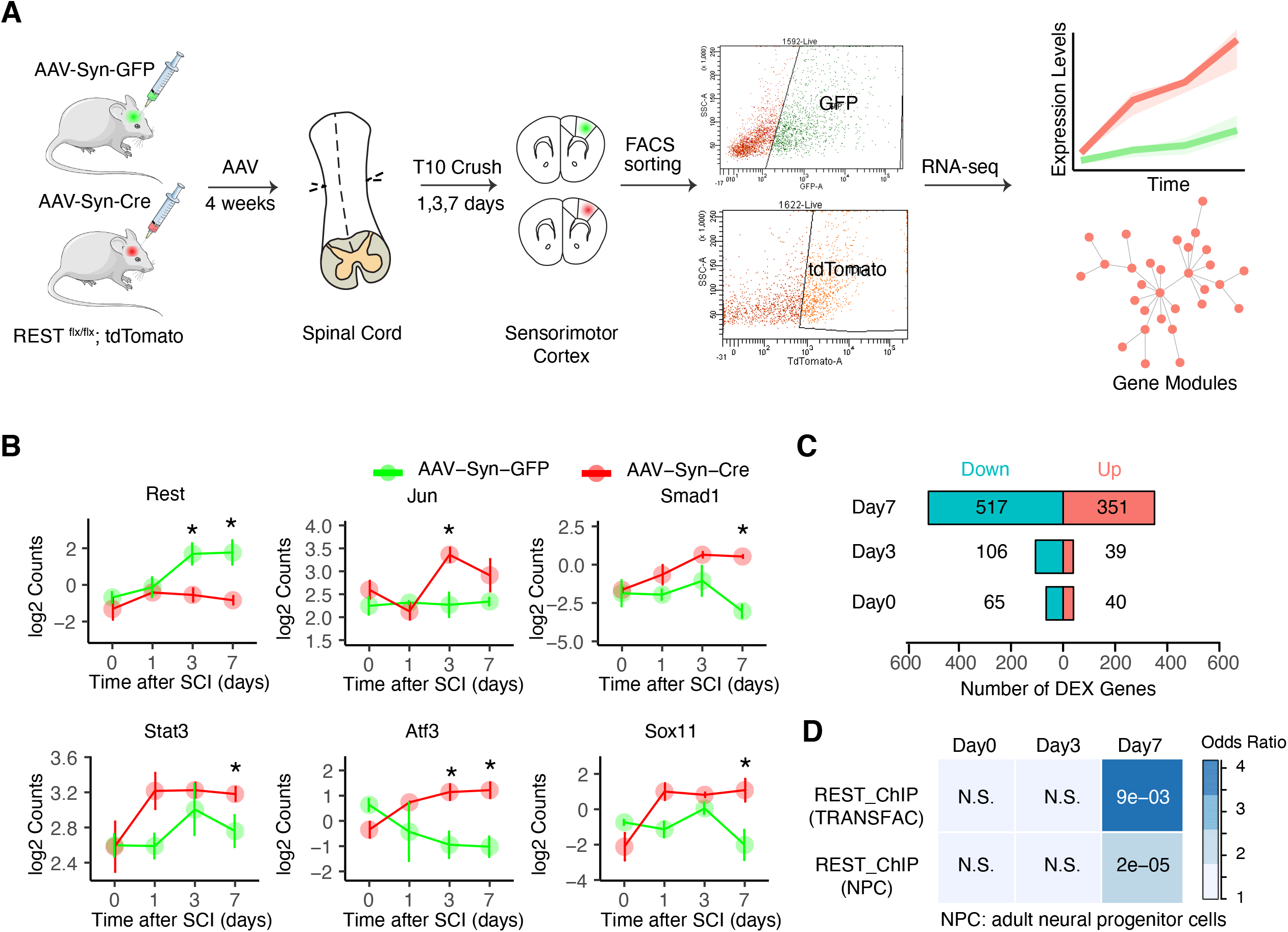
REST deletion in injured cortical neurons enhances expression of regeneration-associated genes and pathways. **(A)** Overview of transcriptional profiling of FACS-sorted corticospinal neurons after SCI. REST*^flx/flx^*; STOP*^flx/flx^*TdTomato mice were injected into the sensorimotor cortex with AAV expressing GFP or Cre recombinase under human synapsin promoter (AAV-Syn-GFP or AAV-Syn-CRE) in order to induce REST deletion and fluorescent labeling of CST projection neurons. Four weeks later, a complete crush injury at thoracic spinal cord level 10 (T10) was performed, followed by FACS sorting and RNA-Seq of GFP or tdTomato-expressing cortical neurons in sham-treated (day 0) and at 1, 3, and 7 days after SCI. n = 3 - 4 mice in each condition. We analyzed transcriptional differences in response to SCI and REST depletion at both individual gene expression level and co-expression network level. **(B)** Expression levels of *Jun, Smad1, Sox11, Stat3, Atf3, and Rest*. Values are mean log2 Counts ± SEM and *p < 0.05 compared to AAV-Syn-GFP at each time point. **(C)** Number of DEGs with FDR corrected p-value < 0.1 and |log2 FC| > 0.3 at each condition. Up-regulated: red; Down-regulated: blue. **(D)** Overlap between up-regulated genes and REST target genes identified from TRANSFAC, the most extensive collection of experimentally determined TF binding sites, or REST ChIP-seq in neural progenitor cells (Mukherjee et al., 2016). Colors indicate odds ratio and values represent p-values (Fisher’s exact test).

We first examined differentially expressed genes in response to injury alone (Figure S2A-B). In wild-type neurons expressing AAV-GFP, SCI up-regulated genes involved in both injury- and regeneration-associated processes at day 1, including apoptosis, neuron projection, cell adhesion, and axon extension (Figure S2C) (*40, 41*). At days 3 and 7 post-injury, however, the up-regulated genes were predominantly associated with injury-relevant pathways involved in oxidative stress, and receptors or channels that increase neural excitability (Figure S2C) (*40, 41*). REST expression levels were increased in sensorimotor cortex neurons at 3 and 7 days post-injury (Figure 3B, AAV-Syn-GFP) in parallel with the expression of injury-relevant gene expression patterns.

The timing of REST expression subsequent to the early, but aborted regeneration pathways, and prior to more subacute injury-related pathways, was consistent with REST potentially repressing regeneration-associated genes and pathways. To test this hypothesis, we compared gene expression responses in sorted, purified, sensorimotor cortex neurons with or without REST deletion at multiple time points post SCI. At early time points following injury, only a few genes were responsive to REST deletion, whereas far more DEGs were identified at 7 days following injury (Figure 3C), consistent with the observed time-dependent increase of REST following SCI. A gene ontology analysis showed that up-regulated genes resulting from REST deletion are involved in regulation of neural transmission, neuron projection, and neurite growth or patterning, while the down-regulated genes are associated with protein translation, mRNA processing, and cell cycle (Figure S3A). Remarkably, expression levels of the core five peripheral axon regeneration-associated TF network genes (*Jun, Smad1, Sox11, Stat3, and Atf3*) (Figure 2) were all up-regulated in REST-depleted neurons (Figure 3B), with *Jun* and *Atf3* significantly increased at day 3 post SCI, and *Smad1, Sox11, Stat3* significantly increased by day 7. Notably, other TFs or known genes in the RAG program that we previously characterized in the PNS (*20*) were also increased by REST depletion (Figure S3B), including immediate early genes induced by peripheral injury (*Egr1)* (*70*), growth-associated proteins (*Gap43, Cap23)* (*71*) (*72*), molecules involved in vesicle and cytoskeletal transport (*Vav2, Syt4*) (*73*) (*74*), cell proliferation (*Pcna*) (*75*), cAMP signaling (*Rapgef4/Epac2*) (*76*) and p38 MAPK signaling (*Atf2, MApkapk2*) (*77*).

REST binds to more than 1300 RE1 sites in the genomes of humans and other mammals (*78*), and the binding of REST to its targets is often context-specific (*79*). To next investigate whether the DEGs are likely to be directly or indirectly regulated by REST in the context of SCI, we compared genes identified in our RNA-seq dataset to genes with experimentally-proven REST-binding genes (*80*), and to a previously published REST ChIP-seq dataset from adult neural progenitor cells (*81*). We found a significant overlap between canonical REST targets and genes up-regulated by REST deletion at day 7 post SCI, though not at other times (Figure 3D; OR _(TRANSFAC)_ = 3.54, OR _(NPC)_ = 2.18; p _(TRANSFAC)_ = 9e-03, p _(NPC)_ = 2e-05, Fisher’s exact test). GO analysis indicates that these overlapping genes are implicated in neural transmission and neuron projection, processes up-regulated by REST depletion (Figure S3A). Overall, these findings indicate that REST is up-regulated by CNS injury (Figure 3B) and that it transcriptionally represses its canonical neuronal target genes, as well as the regeneration-associated TFs, as was predicted by our bio-informatic analysis (Figure 2 and Figure S1). Although the over-representation of REST binding sites with the promotors of RAGs suggested that these effects were mainly direct, it is also possible that REST inhibition improves regeneration by abrogating the transcription of other TFs that are known to limit CNS regeneration, such as *Pten, Socs3, and Klf4*. We did not observe a significant change in expression of these well-defined repressors of regeneration, however (Figure S3C). These findings, and the over-representation of REST binding sites within RAGs, suggest that REST is likely acting independently of these known repressive molecules to regulate axon regeneration.

### REST deletion enhances a co-expression network associated with regeneration

Next, we used Weighted Gene Co-expression Network Analysis (WGCNA) (*69, 82, 83*) to identify network-level changes regulated by REST. Compared with ARACNe, which estimates statistical direct interactions based on mutual information (*48*) and is especially suited for TF-network analysis (Figure 2), WGCNA identifies modules of highly co-expressed genes, with direct or indirect interactions and shared biological functions and pathways (*83*). In addition, we previously showed that WGCNA modules could be further integrated with experimentally validated protein-protein interactions (PPI) to identify protein-level signaling pathways represented by gene networks (*20*). This would not only provide independent validation of the relationship inferred by RNA co-expression, but also important PPI pathways as potential therapeutic intervention.

We performed WGCNA on the RNA-seq data generated from sensorimotor cortex neurons expressing AAV-Syn-GFP (wild-type) or AAV-Syn-Cre (REST-depleted) collected at 1, 3, and 5 days following SCI (Methods; Figure S4 A-C). Based on the correlation of the first principle component of a module, called the module eigengene, with time-dependent changes after injury, we found five modules significantly altered by REST deletion: RESTUP1, RESTUP2, and RESTUP3, which were up-regulated by REST deletion, and RESTDOWN1 and RESTDOWN2, which were down-regulated (Figure 4A lower panel, Figure 4B, Figure S4D). To determine which of these gene modules altered by REST deletion are associated with regeneration, we performed an enrichment analysis between each module and the core RAG co-expression module, which we previously identified to be activated during peripheral nerve regeneration and enriched for regeneration-associated pathways in multiple independent data sets (*20*). This analysis found that the up-regulated module RESTUP1 and RESTUP3 significantly overlapped with a core RAG co-expression module (Figure 4A upper panel).

**Figure 4.**
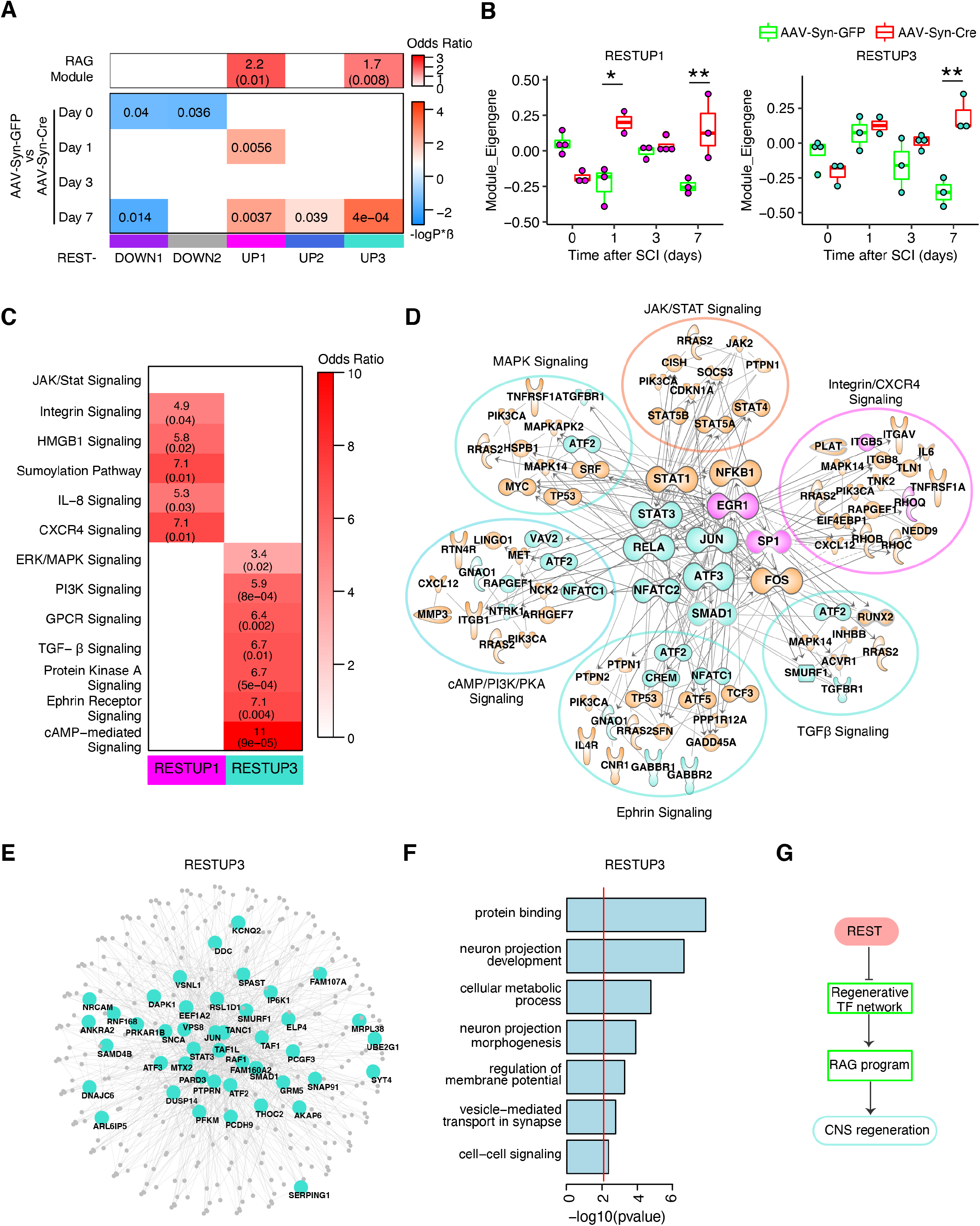
Co-expression network analysis in wild-type and REST-deleted cortical neurons following SCI. WGCNA was performed in REST*^flx/flx^* cortical neurons expressing AAV-Syn-GFP (wild-type) or AAV-Syn-CRE (REST-depleted) in the sham condition (day 0) and at 1, 3, and 7 days after SCI. **(A)** Correlation between module eigengene, the first principle component driving the expression changes of a module, with treatments (bottom panel) and over-representation (hypergeometric test) of regeneration-associated genes (RAGs) (Chandran et al., 2016) within each module (upper panel). The correlation analysis was to identify network-level changes regulated by REST based on the significant module-treatment relationships (Methods). In the correlation heatmap (bottom panel), colors indicate –sign(correlation coefficient)*(log10 p-value). Red indicates a positive correlation and blue indicates a negative correlation. Numbers shown are Bonferroni-corrected p-values. The over-representation analysis was to determine whether modules regulated by REST are enriched with known RAGs activated by peripheral injury. In the enrichment heatmap (upper panel), numbers shown are odds ratio indicating the possibility of enrichment, with hypergeometric p-value in parenthesis. Only modules with significant correlations with REST depletion are displayed in the plot. **(B)** Trajectory of the RESTUP1 and RESTUP3 module eigengenes across different time points after SCI in AAV-Syn-GFP (green) and AAV-Syn-CRE expressing (red) neurons. These two modules are significantly associated with the RAG module activated by peripheral injury. Asterisks denote statistical significance assessed by ANOVA model with Tukey’s post-hoc test: *p < 0.05, **p < 0.01 comparing AAV-Syn-CRE to AAV-Syn-GFP. **(C)** Over-representation (hypergeometric test) of regeneration-associated pathways in RESTUP1 (magenta) and RESTUP3 (turquoise). These regeneration-associated signaling pathways were derived from GO analysis of the RAG module. **(D)** Overlap between protein-protein interactions (PPI) represented by genes in the RESTUP1 and RESTUP3 modules and PPIs from the RAG module. PPIs with significant enrichment for the regeneration-associated pathways are displayed, with the core transcription factors in the center. Each node represents a molecule from the RAG module, colored by orange, while edge represents an experiment-supported PPI between two nodes. Directed edges with arrow represent physical TF-target binding interactions supported by ChIP-datasets from ENCODE and previously published ChIP-ChIP and ChIP-seq experiments. Magenta-colored nodes indicate these molecules also appear in RESTUP1 module, and turquoise indicating molecules also from RESTUP3 module. **(E)** PPI network of RESTUP3 module. The top 70 hub genes which represent the most central genes in the RESTUP3 module were labeled in the network plot. **(F)** GO terms associated with RESTUP3 module. **(G)** A hypothetical model of how REST acts on CNS axon regeneration.

Among the pathways associated with this core RAG module, the RESTUP3 module was enriched with cAMP-mediated, Ephrin-, PKA-, TGFβ-, GPCR- and MAPK signaling, while the RESTUP1 module was modestly enriched with integrin-, chemokine-, and HMGB1 signaling pathways (Figure 4C). To extend this analysis to the protein level, we evaluated the overlap between PPIs from co-expressed genes in RESTUP1 or RESTUP3 and the regeneration-associated PPIs from the RAG module. We found that PPIs from RESTUP3 and RESTUP1 were enriched for very similar regeneration-associated pathways shown by gene-level overlap analysis (Figure 4C), which are linked by members of the core TF regulatory network activated in the regenerating PNS (Figure 4D, Supplemental Table 2), including Jun, SMAD1, STAT3 and ATF3 (Figure 2B, Figure S1A). These core regenerative TFs also appear as module hubs in the PPI network of the RESTUP3 module (Figure 4E). Further GO analysis of general biological pathways represented by these modules showed that the RESTUP3 module is associated with neuronal projection, metabolism, or synaptic transmission (Figure 4F). These analyses support a model whereby inhibition of REST activates a core molecular program driven by a tightly controlled TF network similar to the one activated during peripheral nerve regeneration, along with other complementary pathways, to enable subsequent regenerative processes (Figure 4G).

### REST is a transcriptional repressor negatively correlated with the regeneration state of retinal ganglion cells

To assess the potential generalizability of the bio-informatic predictions derived from spinal cord and peripheral nerve injury above, we extended the same TF regulatory network analysis to another CNS neuronal population, injured retinal ganglion cells (RGCs). RGCs extend axons through the optic nerve, conveying diverse visual features to the lateral geniculate nucleus, superior colliculus, and other relay centers in the di- and mesencephalon, and are a well-established example of CNS neurons that normally exhibit little or no regeneration (*1*); mature RGCs fail to regenerate their axons beyond the site of optic nerve injury and soon begin to die (*84*). However, varying degrees of regeneration can be induced by treatments that include growth factors associated with intraocular inflammation (*85–88*), CNTF gene therapy (*89*), deletion of cell-intrinsic suppressors of axon growth, of which PTEN deletion is the single most effective (*14, 17, 22, 25, 90, 91*), zinc chelation (*92*), physiological activity (*92, 93*), chemical activation of the regenerative gene program (*20*) and, most effectively, by combining two or more of these treatments (*17, 20, 94, 95*).

From our initial bio-informatic predictions comparing PNS and CNS injured tissues, we hypothesized that the disrupted TF network in the injured, non-growing RGCs, similar to the CNS-injured spinal cord tissues (Figure 2D), would re-gain substantial connectivity in RGCs treated so as to be in a more regenerative state. Using mice that express cyan-fluorescent protein (CFP) in RGCs (*96*), we induced robust axon regeneration by combining a strong genetic pro-regenerative manipulation, RGC-selective PTEN knock-down (AAV2-shPten.mCherry; Methods; (*14, 97*)), with intraocular injection of the neutrophil-derived growth factor oncomodulin (Ocm: (*86, 87*) and the non-hydrolyzable, membrane-permeable cAMP analog CPT-cAMP (a co-factor of Ocm) immediately after nerve injury. This combination provides one of the strongest regenerative responses described to date (Figure 5A), while avoiding complications that might be introduced by inducing intraocular inflammation (*15, 16, 87*). Controls received an intraocular injection of AAV2 expressing shLuciferase.mCherry 2 weeks before surgery and saline immediately afterwards. These mice did not exhibit axon regeneration (Figure 5A-B; see Methods). We dissected retinas and FACS-sorted RGCs from non-regenerating, control treatment, or from RGCs exposed to the pro-regenerative combinatorial treatment 1, 3 or 5 days after optic nerve crush injury, followed by transcriptomic analysis via RNA-seq in 8-10 biological replicates for each condition (Figure 5C; Methods).

**Figure 5.**
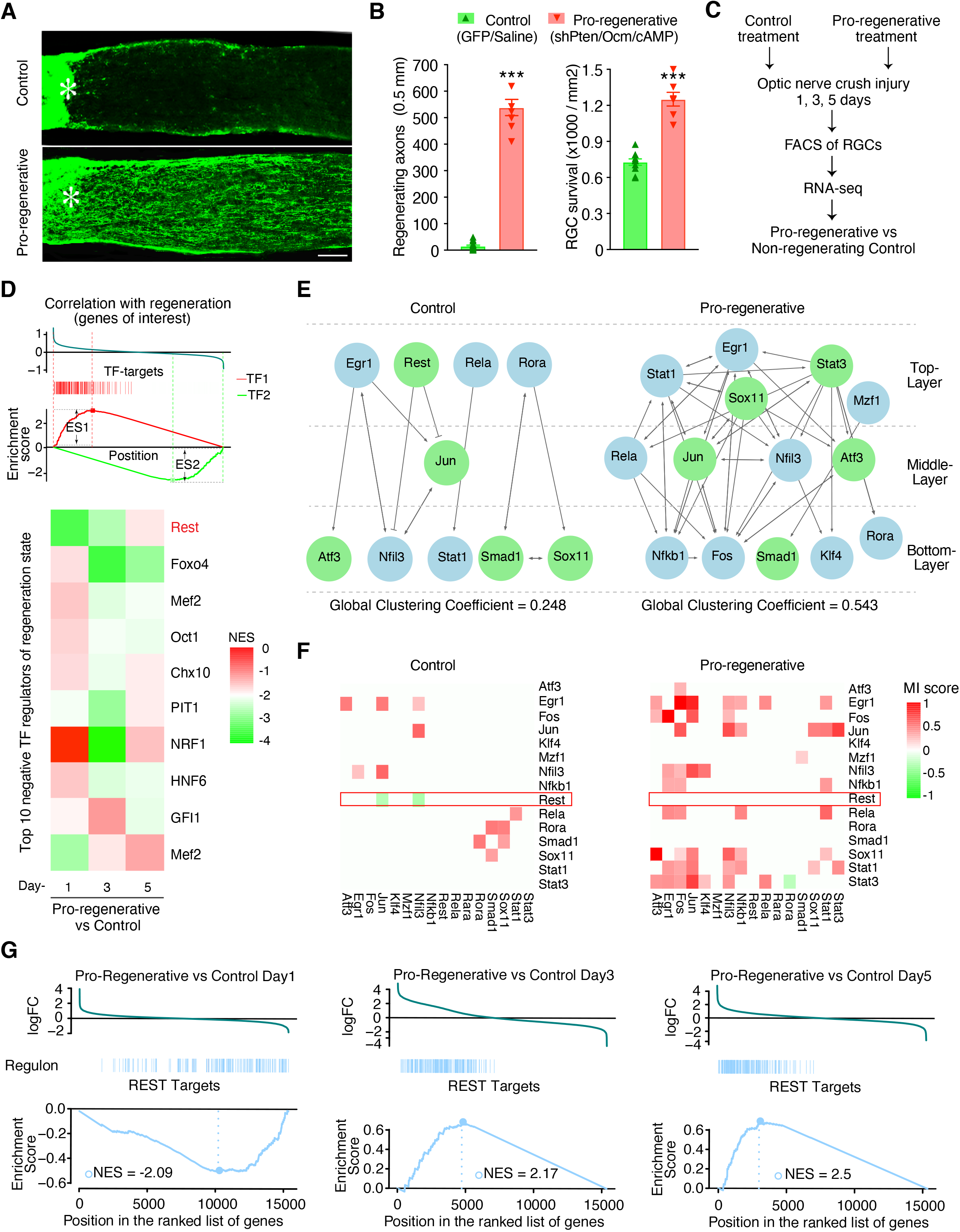
REST is a transcriptional repressor negatively correlated with CNS regenerative state. **(A)** Longitudinal sections through the mature mouse optic nerve show regenerating axons immunostained for GAP43 (green) two weeks after optic nerve crush. Wild-type 129S1 mice expressing cyan-fluorescent protein (CFP) in RGCs received adeno-associated viruses expressing an shRNA to knock down expression of *pten* (AAV2-H1-shPten.mCherry-WPRE-bGHpA, abbreviated: AAV2-shPten.mCherry) as part of the pro-regenerative treatment, or a control virus expressing shLuciferase.mCherry (AAV2-H1-shLuc.mCherry-WPRE-bGHpA, abbr.: AAV2-shLuc.mCherry). After allowing 2 weeks for expression of transgenes, optic nerves were crushed < 0.5 mm distal to the eye and either recombinant oncomodulin plus CPT-cAMP (Ocm+cAMP, other part of pro-regenerative treatment) or saline (control) was injected intraocularly. **(B)** Quantitation of axon growth (left) and retinal ganglion cell (RGC) survival (right). *Asterisk* in A: nerve injury site. Scale bar in A: 120 µm. *** *P* < 0.001, student t-test. **(C)** Schematic depiction of experimental procedures used to generate RNA-seq data from injured RGCs with pro-regenerative treatments or non-regenerating control. B6.Cg-Tg(Thy1-CFP)23Jrs/J mice expressing cyan fluorescent protein in RGCs received the same pro-regenerative or control treatments as in (**A**-**B).** Retinas were dissected and dissociated at 1, 3, or 5 days after surgery, and CFP^+^mCherry^+^ RGCs were separated by FACS. Transcriptomes were evaluated by RNA-Seq to identify transcriptional changes associated with axon regeneration. **(D)** Gene set enrichment analysis (GSEA) to screen TFs correlating with RGC regenerative state. Upper panel: schema demonstrating the principle of GSEA. In this analysis, genes of interest are ranked by their correlations of expression changes with treatments measured by directional p-value, which is calculated as -sign(log Treatment/Control)*(log10 p-value). A positive correlation indicates up-regulation of a gene by pro-regenerative treatment, while a negative correlation indicates down-regulation. Given an *a priori* gene set known to be targeted by a TF, the goal of GSEA is to determine whether this TF’s targets are randomly distributed throughout genes of interest, or primarily found at the top or bottom. An enrichment at the bottom suggests that the TF down-regulates genes of interest, and is thus a negative regulator of the regenerative state (ES <0; TF2 as an example), while an enrichment at the top suggests this TF is a positive regulator of regeneration (ES >0; TF1 as an example). Bottom panel: A total of 1137 TF targeted gene sets were screened and the top 10 negative TF regulators of RGCs’ regeneration state were shown in the heatmap by their normalized enrichment scores (NES). **(E)** Transcriptional regulatory networks comparing RGCs in non-regenerating (control) and regenerating state (pro-regenerative). The networks were constructed using the unbiased, step-wise pipeline described in Figure 2A. **(F)** MI scores of each TF-pair in the networks **(E)** indicating the degree of their correlation. **(G)** Distribution of REST-repressed target genes defined by ARACNe throughout the de-regulated genes by pro-regenerative treatments ranked by log2-fold changes (logFC, pro-regenerative vs non-regenerating) at indicated times following optic nerve crush.

To quantitatively determine a TFs’ association with RGCs’ regeneration state, we first performed gene set enrichment analysis (GSEA) to compare a gene expression signature correlated with the RGC axon regenerative state against ‘tag gene sets’ with known binding sites for TFs (*98*). GSEA returns an enrichment score (ES) of this comparison to determine whether the gene set represented by regeneration-associated genes is enriched in targets of any TFs and if it is a positive or negative regulator of the genes associated with regeneration phenotype (Figure 5D; (*99*). Among the ∼1000 TF-target gene sets unbiasedly tested, REST is ranked as the top negative regulator of the RGC regeneration state-associated gene set at day 1 following injury, which is attenuated on days 3 and 5 after injury (Figure 5D), consistent with REST being an early, upstream event in the regulatory cascade.

We next performed a complementary analysis using the same ARACNe-based pipeline as used in our initial analysis of published PNS and CNS microarray datasets to construct a data-driven, unsupervised, hierarchical network of the regenerative TFs within this new RNA-seq dataset. Similar to CNS injured tissues in the first analysis (Figure 2D), non-regenerative RGCs with control treatment adopt a simpler, less inter-connected, and less structured TF network. This unsupervised analysis showed again that REST appears at the top-layer of the non-regenerating network (Figure 5E, Control), and is negatively correlated with other lower-layer TFs (Figure 5F, Control). By contrast, pro-regenerative treatments re-established a more complex, multi-layered network with higher connectivity (Figure 5E, global clustering coefficient in Control = 0.25, versus the pro-regenerative treatment = 0.54), in which REST is dissociated and the key regenerative TFs (ATF3, Jun, Sox11, Stat3) are more connected (Figure 5F), similar to the microarray data from PNS (Figure 2). Other commonly used statistics for network connectivity such as local clustering coefficient, betweenness centrality, and in- and out-degree (Methods), further revealed significantly higher connectivity for the RAG TFs in the regenerating versus non-regenerating group (Figure S5A). These results from independent datasets and different tissues further support our original bio-informatic predictions that neurons displaying regenerative potential are associated with a highly inter-connected, structured TF-regulatory network. Further, these analyses (*e.g*., Figure 2 and 5) show that REST appears as an inhibitory TF at the apex of a dismantled TF network in the non-regenerating CNS neurons, but is not associated with the highly interacting TF network present in neurons in a regenerating state.

These multiple analyses of independent data suggested that REST is an upstream transcriptional repressor potentially limiting the interactions between lower-level TFs and the expression of regeneration-associated genes. One prediction of this model is that REST target genes should be enriched in RAGs and RAG-associated processes, parallel with GSEA (Figure 5F). We observed 630 transcriptional interactions with REST predicted by ARACNe, including 339 positively regulated (activated) genes and 321 negatively regulated (repressed) genes (Figure S5B, Supplemental Table 4; Methods). Enriched GO terms for genes predicted to be activated by REST include metabolic processes, response to endoplasmic reticulum (ER) stress, and RNA binding and transport (Figure S5C), whereas genes predicted to be repressed by REST are indeed implicated in processes or pathways associated with axon regeneration (*18*), including calcium ion transport, axon guidance, synaptogenesis, CREB- and cAMP-mediated signaling (Figure S5C). The REST-repressed, regeneration-associated gene set was enriched with down-regulated genes at early stages (day 1), which were up-regulated in the later stages of regeneration (day 3 and 5) (Figure 5G, GSEA), suggesting a release of the transcriptional brake by REST on these genes. Altogether, two independent analyses of data from different sources that were focused on identifying key upstream TFs regulating CNS regeneration using unsupervised methods revealed REST to be a key transcriptional upstream repressor of a RAG program, suggesting that it would be a potential novel suppressor of regeneration. Conversely, since REST is a repressor of a pro-regenerative program, these analyses predict that counteracting REST would enhance regeneration after injury. To formally test this model, we next performed several experiments both *in vitro* and *in vivo*, using dorsal root ganglia (DRGs) cultured on a growth-suppressive substrate to model CNS-injured environment and two different *in vivo* models of CNS injury – complete spinal cord injury (SCI) and optic nerve crush.

### REST deletion facilitates, and over-expression inhibits, neurite growth *in vitro*

We first tested the consequences of gain- and loss-of function of REST in dissociated adult dorsal root ganglion (DRG) neurons *in vitro*. We hypothesized that if REST were indeed inhibitory, its depletion should be permissive, whereas its over-expression would inhibit the normal ability of PNS neurons to extend processes. REST depletion was achieved by infecting DRG neurons obtained from REST^flx/flx^; STOP^flx/flx^TdTomato mice (Methods) with adeno-associated virus expressing Cre recombinase (AAV-Cre; Methods). Cells infected with AAV-Cre, but not control virus (AAV-GFP), showed reduced REST mRNA and protein levels with tdTomato expression turned on (Figure S6A-C).

To test the role of REST in a growth-suppressive environment to mimic the injured CNS, we grew DRG neurons on chondroitin sulphate proteoglycans (CSPG), a class of growth-suppressive extracellular matrix molecules present in injured CNS tissue (*4, 100*), and compared this with growth on laminin, a growth-permissive molecule that positively supports extension of injured peripheral axons (*101*). We first determined a CSPG dose that inhibits neurite growth without affecting cell survival (Figure S6D; Methods) and used this concentration to test the effects of REST depletion in DRG neurons. In agreement with previous findings (*102, 103*), DRG neurons treated with AAV-GFP had limited neurite extension when cultured on CSPG (Figure 6A-B). However, REST reduction induced by AAV-Cre (Figure 6C) enhanced neurite outgrowth by ∼40% compared with control neurons (Figure 6A-B, CSPG group), showing that inhibition of REST enables neurite extension of regeneration-competent neurons in a growth-suppressive environment. Notably, REST deletion did not affect neurite extension of DRG neurons when cultured on laminin (Figure 6A-B, laminin group), suggesting that REST-mediated inhibition of growth processes may be activated by a growth-suppressive environment that mimics the injured CNS, such as the presence of CSPG, and is not present in the presence of permissive substrates that support peripheral axonal growth.

**Figure 6.**
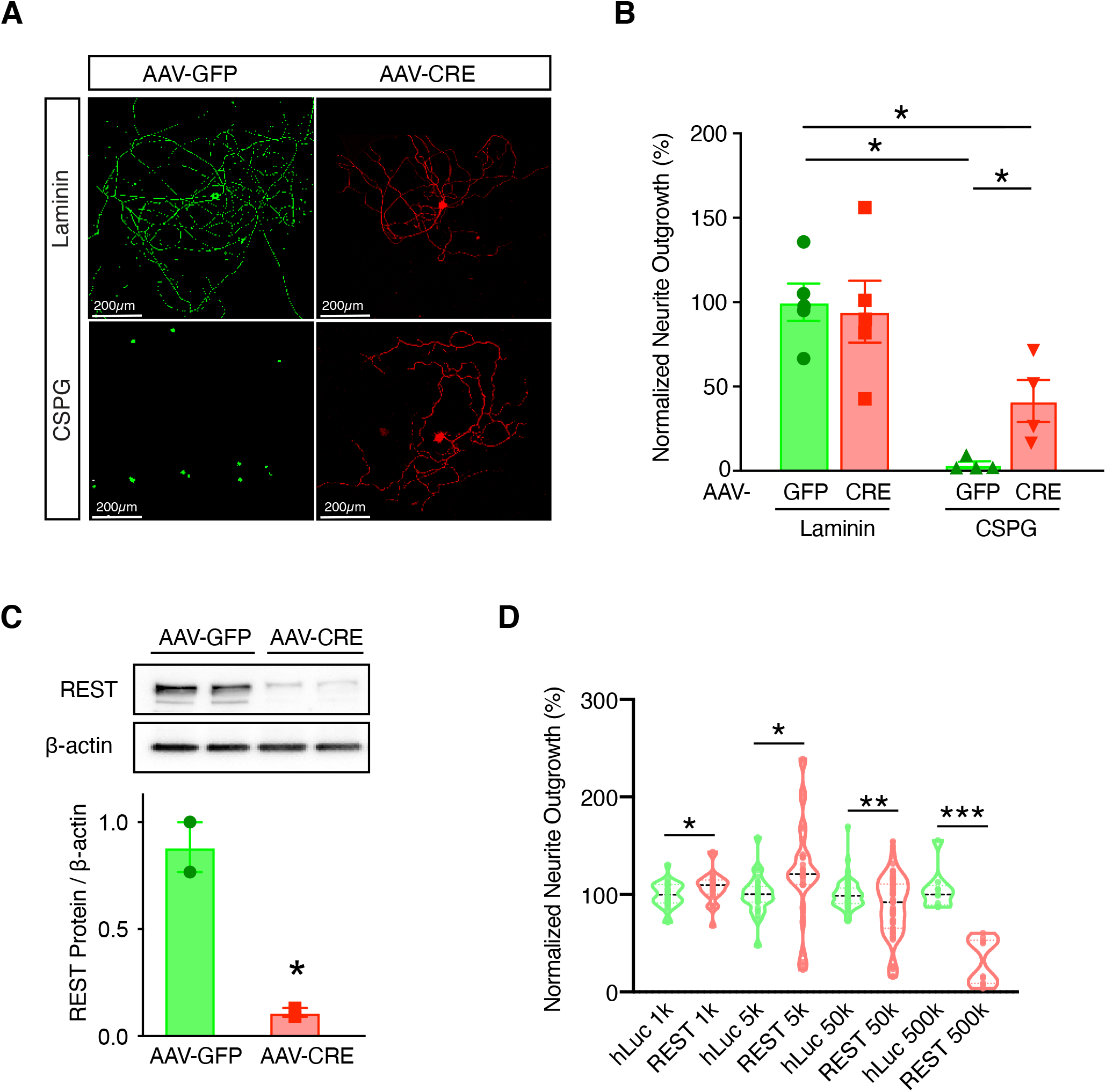
REST inhibits neurite growth *in vitro.* **(A)** Tuj1 (βIII tubulin) staining of REST *^flx/flx^*;tdTomato DRG neurons cultured on CSPG (5 µg/ml) or laminin only (2 µg/ml) and transduced with AAV-GFP (green) or AAV-CRE (red) at ∼ 100,000 genome copies per cell for 7 days to allow the expression of transgenes. **(B)** Mean neurite outgrowth normalized to AAV-GFP infected neurons cultured on laminin. Bars represent mean ± SEM; Asterisks denote statistical significance assessed by two-way ANOVA with Bonferroni post-hoc test (*p < 0.05). **(C)** Representative western blot and quantitation of REST levels in DRG cells transduced with AAV-GFP or AAV-CRE. **(D)** Volcano plot showing the mean neurite outgrowth of re-plated DRG neurons infected with lentiviral constructs expressing either REST (Lv135-REST) or humanized luciferase protein (Lv135-hLuc) as a control driven by the CMV promoter at indicated genome copies per cell for 7 days. Neurite extension was quantified 24 hr following re-plating. Each dot represents the mean neurite outgrowth from 6 wells from a replicate experiment normalized to control at indicated viral doses. Asterisks denote statistical significance assed by Student’s t-test (* p-value < 0.05; ** p-value < 0.01)

We further hypothesized that REST over-expression might inhibit the ability of DRG neurons to extend processes following a PNS injury. To test this hypothesis, we over-expressed REST in cultured DRG neurons for seven days using lentiviral constructs, followed by re-plating, a process to remove existing DRG neurites *in vitro*. This model recapitulates many biochemical and morphological features of an *in vivo* pre-conditioning peripheral nerve injury (Methods) (*104–106*). The efficiency of REST over-expression was confirmed by qPCR (Figure S6D). We observed that increasing REST construct concentration dose-dependently inhibited neurite extension, particularly at the highest concentration (Figure 6D).

### REST deletion enhances corticospinal tract (CST) regeneration after spinal cord injury

To test the predicted role of REST in CST axon regeneration *in vivo*, we injected AAV-GFP or AAV-Cre into the sensorimotor cortex of adult REST^flx/flx^ mice (*107*), where CST neurons of origin are located. Following sham or T10 SCI, CST axons were traced by injecting the anterograde tracer biotinylated dextran amine (BDA) into the sensorimotor cortex (Figure 7A). At 8 weeks post injury, CST axons in mice receiving AAV-GFP exhibited characteristic dieback from the lesion center, consistent with previous reports (*108, 109*). Conditional deletion of REST led to ∼45% more CST axons proximal to the lesion site (Figure 7B-C), suggesting either a lack of dieback in the axons of REST-deficient neurons or a regrowth of axons after injury.

**Figure 7.**
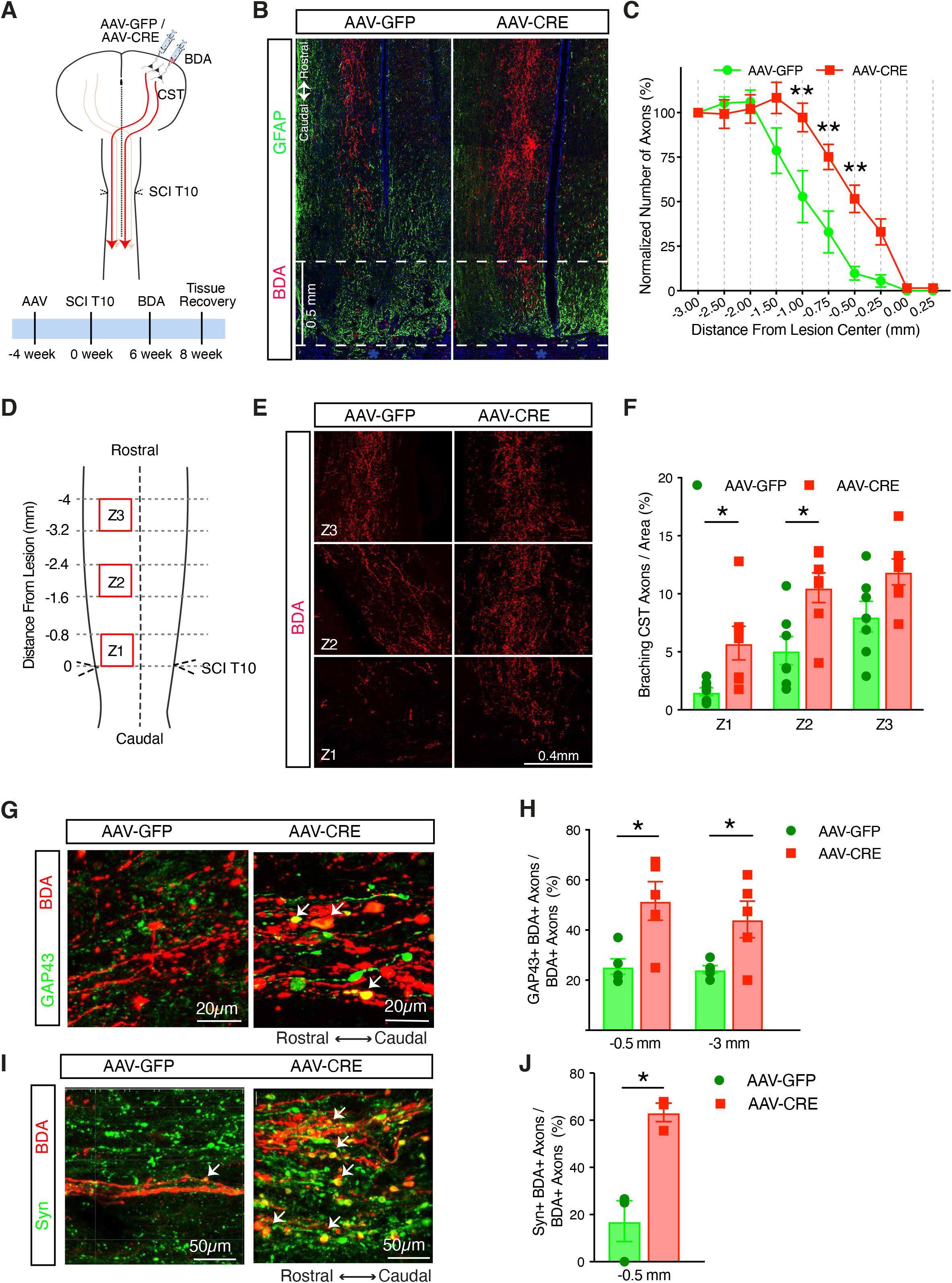
REST deletion enhances corticospinal (CST) axon regeneration after anatomically complete spinal cord crush injury. **(A)** Schematic diagram and timeline of inducing REST deletion and SCI lesions. REST ^flx/flx^ mice were injected into the sensorimotor cortex with AAV2/8.CAG.eGFP.WPRE.polyA (AAV-GFP) or AAV2/8.CAG.Cre-HA.WPRE.polyA (AAV-CRE). Four weeks later, a full crush at thoracic spinal cord level 10 (T10) was performed, followed by cortical injection of BDA to label CST axons. Spinal cords were recovered two weeks after BDA injection. **(B)** Confocal images of BDA-labeled CST axons of lesioned spinal cord also stained for astrocytes (glial fibrillary acidic protein, GFAP). Dashed line represents lesion center (marked with *). **(C)** Intercepts of CST axons with lines drawn at various distances rostral to the lesion center were counted and expressed as percent of the number of intact axons at 3 mm proximally to control for potential variability in the fluorescence intensity among animal. N = 10-12 mice (male and female mixed) in each group; Each dot represents mean ± SEM; **p < 0.01 compared to AAV-GFP at each distance (two-way ANOVA with repeated measures, Bonferroni corrected for multiple comparisons). **(D)** Schematic diagram showing regions along the central canal in horizontal sections of lesioned spinal cord used for quantifying branching of CST axons. Three 0.8 x 0.8 mm^2^ squares (Z1, Z2, Z3) were drawn in the grey matter of each spinal cord as illustrated and the number of axons were counted per square. **(E)** Confocal images of CST axons labeled by BDA in Z1, Z2, and Z3 of each spinal cord. **(F)** Quantitation of the number of axons per area. Bars represent mean ± SEM; Asterisks denote statistical significance assessed by two-way ANOVA with Bonferroni post-hoc test (*p < 0.05 compared to AAV-GFP at each area). **(G-J)** The number of GAP43- or Synaptophysin-expressing axons co-labeled with BDA were counted at 0.5 mm or 3 mm rostral to the SCI crush, and are expressed as percent of BDA labeled axons at respective distances. Confocal images of CST axons (BDA) co-labeled with **(G)** GAP43 or **(I)** Synaptophysin (Syn) at 0.5 mm rostral to the lesion center. **(H)** Quantitation of CST axons expressing GAP43 at 0.5 and 3 mm rostral to lesion center. **(J)** Quantitation of CST axon terminals expressing Syn at 0.5 mm rostral to lesion center. Bars represent mean ± SEM; Asterisks denote statistical significance assessed by one-way ANOVA with Bonferroni post-hoc test **(H)** or Student t-test **(J)** (*p < 0.05 compared to AAV-GFP in each area).

To distinguish between these potential mechanisms, we first examined CST axons 3 days post-injury. Apparent dieback and large numbers of retraction bulbs were observed at this early time point in both control and REST-deleted axons (Figure S7A). We then measured branching of CST axons at 4 weeks post injury which, when increased, is considered to be strong evidence of regenerative growth (*68, 110*) (Figure 7D; Methods). Mice receiving AAV-Cre displayed far more branching from injured CST axons in the area proximal to the lesion center than controls (Figure 7E-F), indicating that REST depletion promotes regenerative axon growth. In addition, REST-deficient CST axons traced by BDA expressed more GAP43 (Figure 7 G-H, GAP43+ BDA+) and synaptophysin (Figure 7 I-J, Syn+ BDA+) than wild-type axons, especially in bouton-like structures in grey matter just proximal to the lesion, indicating the potential of these axons to re-grow and potentially establish pre-synaptic machinery. REST deletion in uninjured mice did not change the number of CST axons (Figure S7B), suggesting that the lack of REST does not affect axon growth in intact or homeostatic states.

### REST inactivation stimulates optic nerve regeneration and RGC neuroprotection

We next tested the role of REST in RGCs, another well-characterized model of CNS regeneration, by intraocular injection of an adeno-associated virus expressing a previously validated dominant-negative REST mutant (AAV2-d/n REST) that includes the DNA-binding domain but lacks the repressor domain of REST (*111*) vs. a control virus (AAV2-GFP: Figure S8A; Methods). After allowing one week for expression of virally encoded d/n REST, we dissected and dissociated retinas and placed the cells in culture (*112*) with or without recombinant oncomodulin, forskolin (to elevate cAMP), and mannose, a necessary co-factor (*87*). Expression of d/n REST caused a modest increase in neurite outgrowth by itself and greatly enhanced levels of neurite outgrowth induced by Ocm/cAMP/mannose (Figure 8A, B). D/N REST also increased RGC survival irrespective of the presence or absence of Ocm/cAMP/mannose (Figure 8A, C).

**Figure 8.**
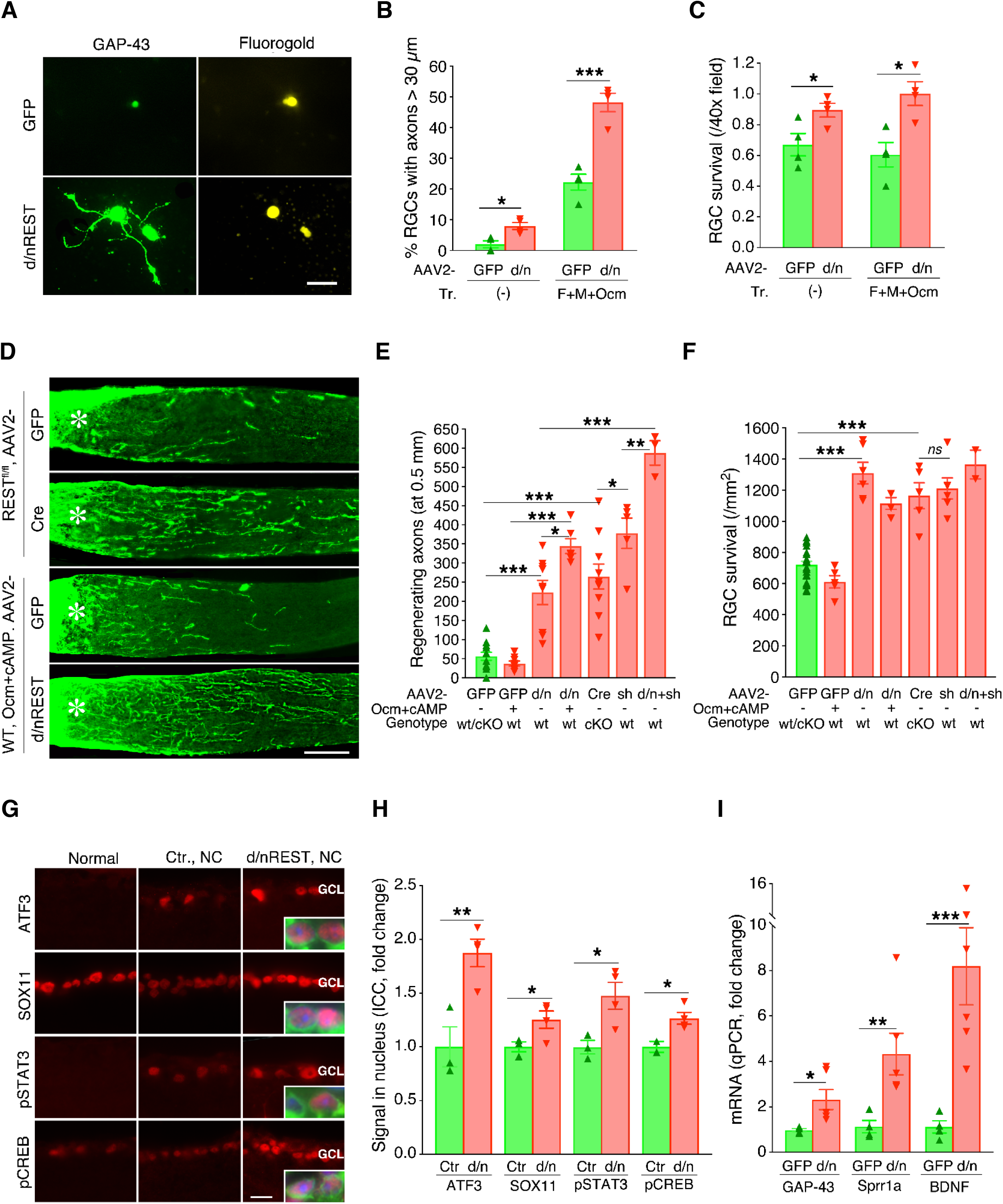
REST inactivation stimulates axon outgrowth from RGCs, optic nerve regeneration, and RGC neuroprotection. **(A-C)** Effect of REST inactivation on adult rat RGCs in culture. Animals received intraocular injections of either AAV2-d/nREST (d/nREST) or AAV2-GFP (GFP) one week prior to dissecting retinas and preparing dissociated cultures. Cells were maintained in the presence or absence of forskolin (to elevate cAMP), mannose, and recombinant oncomodulin (F+M+Ocm) for 3 days. **(A)** GAP-43 immunostaining of RGCs (identified via retrograde labeling with Fluorogold injected into the superior colliculus 7 da earlier). **(B)** Axon outgrowth represented as percentage of RGCs with axons ≥ 30 µm. **(C)** RGC survival in culture. **(D-F)** Effects of REST deletion or antagonism on optic nerve regeneration and RGC survival *in vivo*. REST deletion was obtained by injecting REST^flx/flx^ mice (cKO) intraocularly with AAV2-CAG-Cre.WPREpA (Cre); control REST^flx/flx^ mice received AAV2-CAG-eGFP.WPREpA (GFP). As a second approach, wildtype 129S1 mice (WT) received AAV2-CAG-d/n human REST-HA-SV40pA (d/n) to interfere with REST function or with AAV2-GFP (GFP). In addition to inactivating REST, some WT mice received recombinant Ocm plus CPT-cAMP (Ocm+cAMP). Control mice (green bar in E, F) were pooled from REST^flx/flx^ mice and WT receiving AAV-GFP since there is no difference baseline regeneration between these two groups (Mean±SEM: (71.07±14.65) *vs* (41.57±13.65), *P* = 0.09. Also see Results) **(D)** Longitudinal sections (14 µm) show CTB-labeled axons regenerating through the optic nerve. *Asterisk*: nerve injury site. **(E)** Quantitation of axon regeneration (CTB-positive axons 500 µm distal to the injury site) and **(F)** RGC survival (βIII-tubulin positive cells/mm^2^, average for 8 fields/retina). Both conditional deletion of REST and expression of d/n REST in RGCs increased optic nerve regeneration **(D, E**) and RGC survival **(F)**. **(G-I)** Target gene changes after REST down-regulation. One day after nerve crush, transcription factors predicted to be downstream targets of REST (ATF3, SOX11, pSTAT3), along with pCREB, were elevated in RGC nuclei in mice injected with AAV2-d/nREST (d/nREST, NC) prior to nerve injury (compared to mice receiving the control virus (Ctl. NC) **(G, H)**. Inserts show RGCs at higher magnification: TUJ1: RGCs, *green*; DAPI: nuclei, *blue*; target genes: *red*. Seven days after the nerve crush, mRNAs encoding growth-related proteins were elevated in FACS-selected RGCs expressing d/n REST **(I)**. Statistical tests: **B,C**: student t-test; **E,F**: one-way ANOVA with Bonferroni post-hoc test; **H,I**: multiple t-test. \**P* < 0.05; \*\**P* < 0.01, \*\*\**P* < 0.001. Scale bar in A: 20 µm, in D: 200 µm, in G: 15 µm.

To validate these observations *in vivo*, we used two independent methods to counteract REST (Figure S8A). In the first of these approaches, we examined whether AAV2-d/n REST was sufficient to induce optic nerve regeneration and/or promote RGC survival. Two weeks after optic nerve injury, expression of d/n REST was sufficient to stimulate 43% of the level of axon regeneration (Figure 8D, E) that was achieved with the powerful combinatorial treatment (*pten* deletion, rOcm, CPT-cAMP) subsequently used to generate the transcriptome dataset (c.f. Figure 5A, B). In addition, d/n REST expression more than doubled RGC survival at two weeks post-optic nerve injury (compared to mice injected with AAV2-GFP: Figure 8F), an effect that fully recapitulated the strong neuroprotection afforded by the combination of *pten* deletion, rOcm, and CPT-cAMP (Figure 5B). In parallel to our cell culture studies (Figure 8A-C), we also examined the effect of combining d/n REST expression with Ocm plus cAMP *in vivo*. Whereas a single injection of rOcm + cAMP alone induced little regeneration and no increase in RGC survival relative to untreated controls, combining rOcm + cAMP with the expression of d/n REST increased axon regeneration 55% above the level achieved with d/n REST expression alone (Figure 8D, E). RGC survival was elevated to the same extent as with d/n REST expression alone (Figure 8F).

As an alternative approach (Figure S8A) to investigate the role of REST *in vivo*, we deleted the gene in mature RGCs via AAV2-Cre-driven recombination in mice with homozygous conditional REST alleles and the same TdTomato reporter line (REST^flx/flx^; STOP^flx/flx^ TdTomato) as used in the CST repair studies: see Methods). AAV2-Cre was injected into one eye of REST^flx/flx^; STOP^flx/flx^TdTomato mice, while the contralateral control eye received an injection of AAV2 expressing GFP. REST knock-out was confirmed by tdTomato expression in the retinas of REST^flx/flx^; STOP^flx/flx^TdTomato mice exposed to AAV2-Cre, whereas no TdTomato was observed in control retinas receiving AAV2-GFP. Conditional deletion of REST in RGCs, similar to expression of d/nREST, induced considerable axon regeneration (Figure 8D, E), in this case averaging ∼ 50% of the level induced by the three-way combination of *pten* deletion, rOcm, and CPT-cAMP (Figure 5B). Negative controls were pooled for the different genotypes and viruses used in these studies based on the lack of significant differences in outcomes among controls for AAV2-Cre plus REST^fl/fl^ (strain C57/B6, Mean ± SEM: 71.07 ± 14.65) and for AAV2-d/nREST injections in wild-type 129S1 mice (Mean ± SEM: 41.57 ± 13.65: *P* = 0.09; see legend for Figure 8). In addition, as observed with d/n REST expression, deletion of REST in RGCs doubled the level of RGC survival above that seen in control retinas two weeks after optic nerve injury (Figure 8F), an effect similar to that achieved with the combinatorial treatment used to generate the transcriptional dataset.

Deletion of *pten* is perhaps the most effective single treatment described to date for inducing optic nerve regeneration (*14, 17*). On average, counteracting REST captured ∼ 2/3 of the effect of *pten* deletion on axon regeneration (Figure 8E) and the full effect of *pten* deletion on RGC survival (Figure 8F). Thus, REST can be considered a major suppressor of RGC survival and optic nerve regeneration in mature mice. We also investigated whether *pten* deletion would occlude the effects of counteracting REST, which would suggest that the two share common effector pathways, or whether they might show some degree of additivity. Our results point to partially additive effects on axon regeneration (Figure 8E), suggesting at least some independence of effector pathways.

Accompanying its effects on RGC survival and axon regeneration, expression of d/n REST increased expression of several regenerative TFs (ATF3, SOX11, pSTAT3, pCREB) in the TF regulatory network in RGCs, as assessed by immunostaining retinal sections 1 day after optic nerve injury (Figure 8G, H). At day 7, expression of genes associated with regeneration and/or survival, including *Sprr1a*, *Bdnf* and *Gap-43* were found to be increased based on qPCR using mRNA from FACS-sorted RGCs 7 days after optic nerve injury (Figure 8I: \**P* < 0.05, \*\**P* < 0.01; Methods). These findings are consistent with the elevated expression of key regenerative TFs and effector genes associated with axon growth that we observed in REST-depleted cortical motor neurons, and show that, as with spinal cord injury, REST antagonism enhances central axon regeneration. Thus, we were able to confirm the predicted repressive effects of REST on regeneration based on our systems genomic analysis in two quite distinct models of CNS injury.

## DISCUSSION

We used a stepwise, systems genomics approach to identify upstream transcriptional regulators of intrinsic regeneration-associated gene expression programs in the nervous system. Multiple independent bio-informatic analyses were used to evaluate existing and newly produced gene expression datasets, all of which converged on the transcriptional repressor, REST, as a potential upstream negative regulator of a regenerative gene expression program in the CNS (Figure 1A). We then experimentally demonstrated that disrupting REST activates a core molecular program driven by a tightly controlled TF network similar to the one activated during peripheral regeneration (Figure 1B). This would also predict that counteracting REST would substantially improve regeneration, which was supported in two well established models of CNS injury, the optic nerve and the corticospinal tract (CST) (Figure 1C). These data are consistent with a model whereby REST may act by suppressing the interaction and the expression of pro-regenerative TFs within the RAG network, consistent with its known function as a transcriptional suppressor. Perhaps most importantly, these results firmly demonstrate for the first time that REST represses CNS regeneration *in vivo*, and conversely that its depletion or inhibition by expressing a dominant-negative mutant enhances CNS regeneration.

### TF hierarchies reveal key regeneration-associated factors

Many transcription factors are required to drive growth-associated gene programs for neuronal regeneration (ATF3: (*57, 58*); Jun: (*59, 61*); SMAD1: (*62, 63*); Sox11: (*24, 25, 65, 113*); STAT3: (*21, 60, 89, 114*); KLF family: (*22, 23, 90*)). As this list continues to grow (*19, 20*), efficient strategies are needed to determine how they interact and which TFs are the key factors upstream of regeneration. TF binding is a dynamic process, and a TF can be present or absent from its target loci at different time points and/or under different conditions. In addition, TFs act in a combinatorial manner, forming tiered regulatory networks to drive gene expressions. Therefore, experiments like gain- or loss-of-function of a single or a few TFs at one time is unlikely to recapitulate these TF regulatory events. Here, we used an unsupervised, step-wise bio-informatic approach to characterize the regulatory network structure of regeneration-associated TFs (Figure 2A). By leveraging existing and new gene expression datasets generated in multiple labs and in PNS and CNS injury models at different timescales, we identified a core set of five TFs (Jun, SMAD1, Sox11, STAT3 and ATF3) that occupied a standard, three-tiered core regulatory network (*30, 33, 34*) that was conserved across all PNS datasets (Figure 2C). Each of these core pro-regenerative TFs is increased early after PNS injury (Figure S1A), in agreement with previous findings of their essential role during PNS regeneration (*20, 57–66*) and each connection of TF pairs is experimentally supported (*55, 56*), adding confidence to our bio-informatic predictions.

By contrast, in the non-regenerating CNS (spinal cord and optic nerve), this network loses its three-tiered structure, and instead adopts a simpler, less inter-connected, dismantled structure (spinal cord: Figure 2D; optic nerve: Figure 5E-F). Remarkably, CNS neurons with enhanced regenerative capacity induced by combined genetic and molecular manipulations re-gain the complex, multi-layer TF network with higher inter-connectivity (Figure 5E-F), similar to the TF network induced in the regenerating PNS (Figure 2C). In the dismantled CNS network, REST appears as a top-tier regulator, predicted to inhibit other lower-level TFs. The prediction of REST being a transcriptional repressor was further supported by an independent, unbiased TF-screening approach that evaluated ∼1000 TFs and their experimentally-proven target genes, identifying REST as a top negative regulator of the gene set activated in regenerating CNS neurons (Figure 5D).

Independent analyses of data from different sources that were focused on identifying key upstream TFs regulating CNS regeneration all pointed to REST as a key transcriptional repressor upstream of the core pro-regenerative TFs driving RAG program expression. This prediction was supported by the findings that *Rest* was specifically upregulated across multiple CNS injury datasets (Figure S1A, 3B). When REST is inhibited, Jun, STAT3, Sox11 and ATF3, all members of the core TF regulatory network are up-regulated both in injured cortical neurons (Figure 3B) and in RGCs (Figure 8G-I). Importantly, each of these TFs has been shown independently to promote axonal regeneration, including in the injured CNS in some cases (*19, 21, 22, 24, 25, 61, 66, 113, 114*). These observations, coupled with our data supports a model whereby their up- regulation following REST deletion directly contributes to regenerative growth. Finally, REST depletion enhances gene co-expression programs similar to those activated during peripheral regeneration (*20, 115–117*), involving MAPK-, cAMP-mediated, Neurotrophin- and Integrin signaling pathways driven by the core TFs (Figure 4C-D). Our data suggest a model (Figure 4G) supported by multiple lines of independent data and analyses, whereby REST is induced by CNS injury to suppress the interaction and the expression of several pro-regenerative TFs that act upstream of the RAG program. Thus, inhibition of REST would be expected to release its transcriptional brakes on this program to facilitate axon regeneration in the CNS, which was further validated in multiple experimental models of CNS injury.

### A new role for REST

REST is among the most widely studied transcription factors in the CNS, having been established as a repressor of a large number of genes essential for neuronal function (*78, 118, 119*) and that is predicted to bind and repress close to 2000 putative targets in the mammalian genome. REST and its target genes play important roles in neuronal development as well as the progression of neurological disorders. In the developing nervous system, REST is present in progenitor populations, repressing many genes involved in synaptogenesis, axon pathfinding, and neurotransmission (*78, 79, 118, 120*), but is downregulated at the end-stage of neural differentiation to allow expression of genes that underlie the acquisition of a mature neuronal phenotype (*42–45, 120*). In differentiated neurons, REST is quiescent, but can be activated in response to neuronal insults such as ischemia (*121, 122*) or seizures (*123, 124*), and its expression is linked with neuronal death (*122*). Dysregulation of REST and its target genes has also been associated with the pathogenesis of epilepsy (*125, 126*), Huntington’s Disease (*127*), aging-associated Alzheimer’s Disease (*128*), and decreased longevity (*129*). To date, however, REST has not been linked to CNS repair.

In rodent models of neuropathic pain, REST elevation transcriptionally represses voltage-gated potassium channels in peripheral sensory neurons, resulting in hyper-excitability (*130–132*). Another recent study shows that REST expression transiently increases in response to peripheral injury, but is quickly repressed by an epigenetic regulator, UHRF1, which interacts with DNA methylation enzymes to restrict the transcription of REST, as well as PTEN, a suppressor of cell-intrinsic growth (*133*). We did not observe significant changes of *Rest* expression levels across multiple PNS injury models at different time scales (Figure S1A, PNS1-5). These findings suggest that the expression levels of REST or other intrinsic growth suppressors are tightly controlled in peripherally injured neurons to allow peripheral nerve regeneration.

In view of multiple lines of unbiased bioinformatic data pointing to REST as a novel inhibitor of the intrinsic growth program of CNS neurons, we investigated the effects of REST depletion in two commonly employed models of CNS injury - spinal cord damage and optic nerve injury (*2, 94, 134*). Our data demonstrate for the first time that inhibition of REST enhances regenerative growth in both CNS models, confirming the critical role of REST in suppressing the regenerative competence of CNS neurons. CST axons in animals with REST deletion showed substantially increased growth relative to wild-type controls. We note that although these axons did not grow across an anatomically complete SCI lesion (Figure 7B-C), inability to cross the lesion boundary after complete SCI was expected, as such growth is known to require both intrinsic growth cues and external growth facilitators such as tissue or biomaterial bridges that provide growth-supportive molecules within the lesion site (*8, 135–138*). As a therapeutic strategy for regenerating axons across a complete SCI, it will probably be necessary to augment intrinsic growth capabilities such as REST or PTEN deletion, which activate regeneration-associated genes and pathways in CNS neurons, with an appropriate lesion-bridging substrate (*8, 139*).

In the visual system, expression of a dominant-negative (d/n) REST mutant that retains the DNA-binding domain of the protein but lacks the repressor domain enhanced axon outgrowth in mature RGCs in culture (Figure 8A-C), paralleling earlier observations on the d/n effects of overexpressing RE1 DNA sequences (*140*). In the presence of oncomodulin (Ocm) and a membrane-permeable, non-hydrolyzable cAMP analog, expression of d/n REST led to extraordinary levels of RGC axon outgrowth (Figure 8A-C). *In vivo*, we investigated the role of REST in optic nerve regeneration by two approaches, overexpressing the dominant-negative REST mutant and conditional deletion of the *Rest* gene in RGCs. The effect of counteracting REST was considerable; regeneration induced by counteracting REST was approximately 2/3 of that induced by PTEN deletion, a treatment that provides perhaps the strongest regeneration induced by a single genetic manipulation to date (Park et al., 2008), and roughly half the robust level of axon regeneration induced by *Pten* deletion combined with Ocm and cAMP elevation (Figure 8D-E), the potent combinatorial treatment used to generate our original regeneration RNA Seq dataset. Combining d/n REST expression with Ocm plus CPT-cAMP brought the level of regeneration even closer to that induced by the strong combinatorial treatment, while a combined treatment to knock down PTEN and counteract REST in RGCs led to considerably greater regeneration than either one alone. In addition, expression of d/n REST or REST knock-down was sufficient to double levels of RGC survival, affording the same level of neuroprotection as either combinatorial therapy or PTEN deletion alone, which is notable, since to date, few factors other than PTEN deletion enhance both RGC regeneration and survival. For example, ATF3 is pro-survival but has no effect on RGC regeneration (*141*); Sox11 is pro-regenerative, but when overexpressed, lead to the death of alpha RGCs (*25*); and STAT3 is pro-regenerative, but does not increase survival (*89*).

### Limitations and future directions

Here we demonstrate via several lines of experimental evidence that REST is an inhibitor of CNS axon regeneration. Based on multiple forms of bio-informatic and experimental analyses, we present a model whereby REST acts via repression of pro-regenerative genes, whose regulatory elements it binds. Although we know that REST does repress this regenerative program, and its reduction leads to regeneration, we cannot yet say with certainty that its effects on regeneration are solely via this pathway. Thus, we view this as a working model that warrants further testing. We also note that the genetic manipulations required for direct testing of this model (e.g. simultaneous suppression of multiple core regeneration-associated TFs in the context of REST deletion) are at the very least daunting and at the limit of current experimental tractability. It is also plausible that transcriptional regulation by REST is one of several mechanisms by which its deletion promotes cell-intrinsic growth. From this perspective, it is likely that other key regulators act synergistically with REST to control CNS regeneration. One potential REST-interacting factor could be PTEN, inhibition of which, plus Oncomodulin and cAMP elevation up-regulates a regeneration-associated gene set that is predicted to be repressed by REST (Figure 5G, S5 B-D). As a protein phosphatase, PTEN antagonizes the PI3K-AKT-mTOR pathway to inhibit protein translation, cell cycle progression and cell survival (*63*), as well as transcriptional regulation of cell-growth-associated genes through inhibition (*142–144*). Our findings indicate that REST is likely not acting via eliciting changes in PTEN, or the downstream canonical mTOR pathway to regulate regeneration, as our gene expression data show no change in the levels of *Pten* with REST deletion (Figure S3E), nor do we see changes in phosphorylation of ribosomal protein S6, which would be indicative of changes in the mTOR pathway (Figure S8 B-C). In addition, we observed additive effects of *Pten* deletion combined with counteracting REST, suggesting that the two treatments may activate downstream effector pathways that are at least partially separate (Figure 8E-H). Future studies on how REST interacts with PTEN and other pro-regenerative manipulations will be important in optimizing therapeutic strategies for CNS repair.

Further studies will also be required to clarify the precise molecular mechanisms by which REST acts on the core TF network in the RAG complex to regulate regeneration-associated pathways during CNS repair, and to explore other possible mechanisms. REST may be recruited directly to the regulatory sites for repressing regeneration-associated transcription following CNS injury. ChIP-seq studies have shown that REST can directly bind to regenerative TFs such as Sox11, KLF6, Jun and STAT3 (*79, 81, 145*). Whether REST binds and represses additional regenerative factors in the context of axonal injury needs to be further investigated. It is also possible that REST deploys additional mechanisms of regulating CNS regeneration in addition to acting directly on the core TFs. As a transcriptional regulator, REST can induce chromatin remodeling (*46, 47*), a process that rearranges the chromatin to facilitate or prevent gene transcription. Overall, future studies on a genome-wide profiling of REST occupancy induced by CNS injury or chromatin regulatory changes with and without REST inhibition in CNS neurons will be necessary to identify how REST regulates regeneration-associated transcription to enhance CNS repair. In addition, the mechanisms by which REST itself is regulated in the context of CNS injury is unclear. Others have shown that REST can be regulated post-transcriptionally (*146*), post-translationally via ubiquitination/deubiquitination (*147, 148*), and by cytoplasmic sequestration (*127*). Thus, investigating how REST is regulated in CNS neurons in growth-permissive or non-permissive states may further illuminate non-transcriptional mechanisms underlying CNS regeneration. The unbiased discovery of REST as a regulator of CNS axon regeneration and the validation of REST’s new role across different models of CNS injury provide a proof of concept for the power of our bio-informatic framework as a platform for discovery. In view of the complexity of how REST interacts with the genome, further work will be required to understand more fully how REST regulates the regenerative state of CNS neurons. At the same time, it will be important to investigate the potential of REST manipulation to enhance the ability of other pro-regenerative treatments to improve outcome after CNS injury and to move such treatments towards clinical application.

## Methods

### Animals

Mouse lines, including 129S1, C57BL/6J, loxP-REST-loxP (REST^flx/flx^), B6.Cg-Tg(Thy1-CFP)23Jrs/J, and Rosa26-CAG-loxP-STOP-loxP-tdTomato (STOP^flx/flx^ TdTomato), were purchased from Jackson Laboratory. REST^flx/flx^; tdTomato homozygous mice were generated by crossing REST^flx/flx^ (*107*) and STOP^flx/flx^ TdTomato mice. Young adult mice between 4-6 weeks old including both sexes were used for all experiments in spinal cord studies and 8 -12 week old animals in optic nerve regeneration studies. Experiments performed at University of California, Los Angeles were approved by the Animal Research Committee of the Office for Protection of Research Subjects. Experiments performed at Boston Children’s Hospital were approved by the Institutional Animal Care and Use Committee (IACUC).

### Spinal cord injury and corticospinal tract (CST) injections

Surgical procedures for spinal cord injury and CST injections in mice were similar to those described previously (*8, 108, 109, 149*), and were conducted under general anesthesia with isoflurane using an operating microscope (Zeiss, Oberkochen, Germany), and rodent stereotaxic apparatus (David Kopf, Tujunga, CA). The adeno-associated virus-green fluorescent protein (AAV-GFP) or adeno-associated virus expressing Cre recombinase (AAV-Cre) were obtained from Boston Children’s Hospital Viral Vector Core. The viruses referred to as AAV-GFP and AAV-Cre were AAV2/8.CAG.eGFP.WPRE.polyA and AAV2/8.CAG.Cre-HA.WPRE.polyA, respectively. A total of 2 μl AAV2/8-GFP or AAV2/8-Cre virus at a titer of ∼10^13^ gc /ml was injected into the left cerebral motor cortex at the following coordinates (in mm): anteroposterior/mediolateral: 0.5/1.5, 0.0/1.5, -0.5/1.5, - 1.0/1.5, at a depth of 0.5 mm. Four weeks later, a laminectomy was performed at T10, and the spinal cord was crushed using .1mm-wide customized forceps. To trace corticospinal tract axons, 2 μl biotinylated dextran amine 10,000 (BDA, Invitrogen, 10% wt/vol in sterile saline) was injected at the same coordinates as the AAVs into the left motor cortex six weeks after SCI. Mice that underwent surgical procedures were placed on a warming blanket and received an analgesic before wound closure and every 12 h for 48 h post-injury.

### Immunostaining of spinal cord and cortex

Spinal cords were recovered and stained as previously described (*8, 109*). Following terminal anesthesia by pentobarbital, mice were perfused transcardially with 10% formalin (Sigma). Spinal cords and brains were removed, post-fixed overnight, transferred to buffered 30% sucrose for 48 h, embedded in O.C.T. Compound (Tissue-Tek, Sakura-Finetek/VWR) and cryostat-sectioned at 30 μm. Serial horizontal sections of spinal cord containing the lesion sites and brain containing the viral injection sites were cut and processed for immunostaining. The following primary antibodies were used: GFAP (DAKO, 1:1000, free-floating), GAP43 (1:1000, Benowitz lab), Synaptophysin (Synaptic Systems, 1:1000, free-floating), RFP (1:500, Invitrogen, free-floating), and NeuN (1:500, Millipore, free-floating). BDA tracing was visualized with streptavidin-HRP (1:300, PerkinElmer) antibodies plus Cy3-TSA (1:200, PerkinElmer). Sections were cover-slipped using Prolong Diamond Antifade Mounting media with DAPI (ThermoFisher) to stain cell nuclei.

### Quantitation involving CST axons

To quantify total labeled CST axons, we counted intercepts of BDA-labeled fibers with dorsal-ventral lines drawn at defined distances rostral to the lesion center. Similar lines were drawn and axons counted in the intact axon tract 3 mm proximal to control for potential variability in the fluorescence intensity among animals. Fibers were counted on at least two sections per mouse, and the number of intercepts near or in the lesion was expressed as percent of axons in the intact tract divided by the number of evaluated sections. To quantify the number of branching axons from the main CST, three 0.8 x 0.8 mm^2^ squares (Z1, Z2, Z3) were drawn along the central canal at defined distances rostral to the crush site. The number of axons were counted in each square, and are expressed as percent of area per section for each mouse. The number of GAP43- or Synaptophysin-expressing axons co-labeled with BDA were counted at 0.5 mm and 3 mm rostral to the SCI crush, and are expressed as percent of BDA labeled axons at respective distances. We examined BDA labeling 3 mm caudal to the lesion center to make sure the SCI lesions were complete. All axon counts were carried out by an investigator blind to the identity of the cases.

### Optic nerve crush and intraocular injections

Surgical procedures for optic nerve injury and intraocular injections in mice were similar to those described previously (*15, 16, 86, 112, 150*). To investigate REST functions *in vivo*, we either deleted REST in RGCs or expressed a dominant-negative mutant form of REST (*111*) (d/n REST, gift of Dr. Gail Mandel, OHSU). For the former, REST^flx/flx^-tdTomato mice received an intraocular injection of either AAV2-CAG-Cre.WPREpA (AAV2-Cre, to preferentially delete the gene in RGCs) or, as a control, AAV2-CAG-eGFP.WPREpA (AAV2-GFP). In the latter studies, 129S1 wildtype mice received AAV2-CAG-d/n human REST-HA-SV40pA (AAV2-d/nREST) to inactivate REST function or AAV2-GFP as a control. All viruses were injected in a volume of 3 µl and a titer of 1 x 10^13^ gc/ml 2 weeks prior to optic nerve crush to insure adequate time for gene deletion or transgene expression at the time of nerve damage. Two days prior to the end of a 14-day survival period, cholera toxin B subunit (CTB, 3 µl/eye, 2 µg/µl, List Biological Laboratories, Inc., 103B) was injected intraocularly as an anterograde tracer to label axons regenerating through the optic nerve.

In some studies, 129S1 mice received an intraocular injection of AAV2-d/nREST or an AAV2 control virus two weeks before the optic nerve crush and were euthanized at day 1 or day 7 after nerve injury. Retinas from these mice were prepared for immunostaining of serial sections (details in Methods: *Immunostaining of retinal sections and intensity quantitation*).

To investigate the transcriptome of RGCs during optic nerve regeneration or after counteracting REST, we carried out optic nerve crush surgery with different intraocular treatments *in vivo*, then used FACS to isolate RGCs for subsequent analyses (details in Methods: *FACS isolation of retinal ganglion cells*)

### Quantitation of optic nerve regeneration and RGC survival

Following transcardial perfusion with saline followed by 4% paraformaldehyde (PFA), optic nerves and retinas were dissected out and post-fixed with 4% PFA for 2 hours (RT). Nerves were transferred to 30% sucrose at 4°C overnight before being frozen in O.C.T. Compound (Tissue-Tek, Sakura-Finetek/VWR) and sectioned longitudinally on a cryostat at 14 µm thickness. Regenerating axons were visualized by immunostaining for CTB (1:500, Genway Biotech, GWB-7B96E4) and were quantified in 4-8 sections per case to obtain estimates of the total number of regenerating axons at 0.5 mm distally from the injury site as described (*86, 150*). Whole retinas were immunostained for βIII-tubulin (1:500, free-floating. Abcam) to identify RGCs, and RGC survival was evaluated in 8 pre-designated fields in each retina as described (*86*).

### Immunostaining of retinal sections and quantitation of signals

Animals injected intraocularly with AAV2-d/nREST or a control virus underwent optic nerve crush surgery 14 days later and were euthanized and perfused after another 1 day or 7 days (Methods: *Optic nerve crush and intraocular injections*). Eyes were dissected out, post-fixed for 2 hours at RT, then transferred in 30% sucrose at 4°C overnight. After embedding in O.C.T. and cryostat-sectioned at 14 µm, retinal sections were immunostained with primary antibodies against various proteins, including several transcription factors (anti-ATF3, 1:100, Abcam Ab207434; anti-SOX11, 1:500, Millipore ABN105; anti-pSTAT3, 1:200, Cell Signaling D4769; anti-pCREB, 1:100, Alomone Labs; and anti-βIII tubulin [TUJ1], 1:500, Biolegend to identify RGCs) at 4°C overnight followed by the appropriate fluorescent secondary antibodies the next day. Stained retinal sections were imaged using equal exposure conditions across all sections in both control and treated groups. Staining intensity was measured with Image J software on each individual RGC that was labeled by the TUJ1 antibody, and data were averaged from 50 - 100 consecutively encountered RGCs across 3 different areas from each retina, 3 – 4 retinas per group, and was compared between the control and treatment groups for each antibody.

### Retrograde labeling of RGCs and preparation of dissociated retinal cultures

The procedure for retrograde labeling of RGCs has been described previously (*87, 112*). Briefly, to distinguish RGCs from other cells in dissociated mixed retinal cultures, we injected 2% of Fluorogold (FG, Fluorochrome) into the superior colliculus (SC) bilaterally in adult rats. At the same time, rats received intravitreal injections of either AAV2-d/n REST or AAV2-GFP viruses. After allowing one week for FG transport and viral gene expression in RGCs (*86, 87, 112*), retinas were dissected, dissociated with papain, and the dissociated retinal cells were plated on poly-L-lysine pre-coated culture plates. To obtain a baseline of plated RGCs from different retinas, we carried out an initial quantitation of FG-labeled RGC numbers in culture 5 – 12 hours after plating cells. Axon outgrowth and RGC survival were evaluated after 3 days in culture, and each experimental condition was tested in quadruplicate. Counting was carried out using a fluorescent inverted microscope by an observer who was blind to treatment. RGCs were identified by FG labeling under fluorescent illumination, then evaluated for axon growth using phase-contrast to obtain the percentage of RGCs that extended axons ≥ 30 µm in length. Cell survival is reported as the number of FG-positive RGCs per 40X microscope field averaged over ≥ 30 pre-specified fields per well. The RGC numbers counted at D3 were first normalized by their own initial number at 5 -12 hours after plating, then averaged within the group. In some cases, cultured cells were immunostained with a rabbit monoclonal antibody to GAP-43 (1:500, Abcam, Cat#: ab75810) to visualize regenerating axons.

### Dissociated dorsal root ganglion neuronal cultures and neurite outgrowth assay

Adult C57BL/6J dissociated DRG cells were plated at a concentration of 5,000 – 10,000 cells / ml in tissue culture plates coated with poly-L-lysine (Invitrogen, 0.1 mg/ml) and laminin (Invitrogen, 2 ug/ml) only or with CSPG (Millipore, 5 ug/ml) cultured in Neurobasal A medium (Invitrogen) containing B27 supplement, penicillin, streptomycin, 1 mM L-glutamine, 50 ng/ml NGF, and 10 mM AraC at 37°C. REST overexpression was performed by transducing DRG neurons with lentiviral constructs containing either REST (Lv135-REST) or humanized luciferase protein (Lv135-hLuc) as a control driven by the CMV promoter (GeneCopoeia). DRG neurons were replated 7 days after the viral infection. Replated neurons were allowed to grow for another 17-24 hr before quantifying neurite outgrowth. To test neurite growth on laminin or CSPG, DRG neurons dissected from REST^flx/flx^ mice were dissociated and REST was depleted by infecting neurons with AAV-CRE (experimental) or AAV-GFP (control), the same AAVs used in the Methods section *“Spinal cord injury and corticospinal tract (CST) injections”,* at a viral titer of ∼100,000 genome copies per cell. Neurite growth was measured after 7 days, and each experimental condition was tested in triplicate. To stain DRG neurites, cells were fixed with 4% paraformaldehyde and blocked for one hour at room temperature in PBS with 0.05% Tween-20 + 0.01% Triton-X + 1% BSA + 5% goat serum, followed by primary antibody incubation with ß-III-tubulin (Biolegend, 1:500) overnight at 4 °C in blocking solutions and secondary antibody (Invitrogen, 1:500) for 1-2 hr at room temperature. For quantification of DRG neurites, at least 9 images were randomly taken from each replicate using a Zeiss Confocal Microscope at 20x. Neurites were counted using Imaris Surface Rendering function, and the average neurite surface per neuron was quantified.

### qRT-PCR

RNA from various treatment groups was extracted using the RNeasy kit (Qiagen), reverse-transcribed to cDNA with iScript cDNA Synthesis kit (Bio-Rad) or Quantitect Reverse Transcription kit (Qiagen) for low-input samples. Real-time qPCR was carried out with iTaq Universal (Bio-Rad) or Quantitect (Qiagen) SYBR Green supermix. The primers used in qPCRs were:

SPRR1a F: GTCCATAGCCAAGCCTGAAGA; R: GGCAATGGGACTCATAAGCAG; GAP-43 F: GTTTCCTCTCCTGTCCTGCT; R: CCACACGCACCAGATCAAAA. BDNF F: CACTGTCACCTGCTCTCTAGGGA; R: TTTACAATAGGCTTCTGATGTGG; ATF3 F: CTGGGATTGGTAACCTGGAGTTA; R: TGACAGGCTAGGAATACTGG; REST F: CGACCAGGTAATCGCAGCAG; R: CATGGCCTTAACCAACGACA; 18S F: CGGCTACCACATCCAAGGAA; 18S R: GCTGGAATTACCGCGGCT. Relative expression levels in experimental groups were first normalized to those of the reference gene 18S rRNA, then normalized by the relevant control group depending on the experimental design. Statistical significance among groups was evaluated by one-way ANOVA followed by Bonferroni or Tukey corrections.

### Western blots

Lysates from DRG neurons were run on 4-12% Bis-Tris gradient gels and proteins were transferred to PVDF membranes that were incubated with antibodies to REST (Abcam, 1:1000), using anti-β-actin as a loading control. Quantitation of western blot results was carried out with ImageJ software.

### FACS isolation of adult cortical motor neurons

Surgeries and AAV injections were carried out in the same way as described in the Methods section “*Spinal cord injury and corticospinal tract (CST) injections*”. In order to induce neuron-specific REST depletion, we used AAVs expressing GFP or Cre recombinase under the human synapsin promoter. Adult mouse brain tissue was dissociated as previously described (*151*). Briefly, sensorimotor cortex injected with AAV-Syn-GFP or AAV-Syn-CRE to induce tdTomato expression from REST^flx/flx^; tdTomato mice was immediately dissected into ice-cold Hibernate A without calcium (BrainBits, HA – Ca). Tissue was digested by activated papain (Worthington, resuspended in 5 ml HA-Ca) with 100 μl DNase I (2 mg/ml, Roche) in a 37 °C incubator shaking orbitally for 30 min. Digested tissue was triturated gently until clumps disappeared, spun down, and resuspended in 3 ml HA –Ca containing 10% v/v ovomucoid (Worthington, resuspended in 32 ml HA –Ca). Cell debris was removed using discontinuous density gradient containing 3 ml tissue mixture on top of 5 ml ovomucoid solution. Cells were spun down at 70 x G for 6 min and the pellet was resuspended in 1.8 ml Hibernate A low fluorescence (HA-LF; BrainBits) to create a mononuclear cell suspension. Miltenyi myelin removal kit was used to further reduce the amount of debris according to the manufacturer’s protocol. Briefly, 200 μl myelin removal beads (Miltenyi) were added to the cell suspension and incubated at 4 °C for 15 min, then the cell suspension was centrifuged at 300 x G for 10 min at 4 °C. The pellet was resuspended in 1 ml of HA-LF and applied to LS columns (Miltenyi) attached to MACS magnetic separator in order to remove beads with myelin. Flow-through, as well as two -- 1 ml washes with HA-LF, were collected, centrifuged at 600 x G for 5 min at 4 °C and resuspended in 750 μl HA-LF. Myelin-depleted samples were labeled with live cell marker DRAQ5 (1 μl per sample; Thermo Fisher Scientific) and dead cell marker NucBlue (1 drop per sample; Invitrogen). Samples were FACS-sorted on a Becton Dickinson FACS Aria cell sorter gating for DAPI−/DRAQ5+/GFP+ cells directly collected in 100 μl of RA1 lysis buffer with 2 μl tris(2-carboxyethyl)phosphine (TCEP) from NucleoSpin RNA XS kit (Clontech).

### FACS isolation of retinal ganglion cells

To investigate the transcriptome of RGCs undergoing axon regeneration, B6.Cg-Tg(Thy1-CFP)23Jrs/J mice, which express cyan-fluorescent protein selectively in RGCs (*96*), received intraocular injections of either a well characterized adeno-associated virus expressing shRNA against PTEN mRNA (*97*) and mCherry (AAV2-H1-shPten.mCherry-WPRE-bGHpA, in short: AAV2-shPten.mCherry), or a control virus expressing shLuciferase.mCherry (AAV2-H1-shLuc.mCherry-WPRE-bGHpA, in short: AAV2-shLuc.mCherry). After allowing two weeks for expression of virally encoded genes, mice underwent optic nerve crush. Experimental mice received an intraocular injection of recombinant oncomodulin (rOcm, 90 ng) plus CPT-cAMP (cAMP, 50 µM, total volume = 3 µl); control mice received intraocular saline. At one, three or five days post-surgery, mice were euthanized, retinas were dissected and dissociated by gentle trituration in the presence of papain, and cells were separated by fluorescent-activated cell sorting (FACS, BD Biosciences) on the basis of being positive for both CFP and mCherry (*i.e.*, virally transfected RGCs). We typically obtained 2,000 – 11,000 RGCs per retina and pooled RGCs from 2-3 similarly treated retinas for one sample depending on the number of sorted cells; each condition was repeated at least 8 times in independent experiments.

To investigate the effects of REST manipulations on regeneration-associated TFs and other genes, we injected WT 129S1 mice intravitreally with AAV2-d/nREST (vs. AAV2-GFP in controls) and, at the same time, injected Fluorogold (Fluorochrome) into the superior colliculus (SC) to retrogradely label RGCs. The optic nerve was crushed two weeks later and, after allowing a one week survival period, we euthanized mice, dissected the retinas, dissociated cells (for details see retinal dissociated cell culture) and selected FG-positive RGCs by FACS. RNA from sorted RGCs was extracted for each sample and prepared for real-time qPCR analysis.

### Transcriptional regulatory network analysis

A stepwise pipeline was used to construct a hierarchical TF network from gene expression datasets. Step 1: The Algorithm for Reconstruction of Accurate Cellular Networks (ARACNe) (*48*) was applied to each of the gene expression profiling datasets to infer directionality among TFs using RTN package (*152*). Pair-wise mutual information (MI) scores were computed and non-significant associations were removed by permutation analysis (permutation = 100; FDR adjusted p value < 0.05; consensus score = 95%). Unstable interactions were removed by bootstrapping, and indirect interactions such as two genes connected by intermediate steps were removed by data-processing inequality (DPI) of the ARACNe algorithm. Step 2: To further confirm the directionality inferred by ARACNe, we examined evidence of physical TF-target binding observed by multiple ChIP-Seq or ChIP-ChIP databases (*30, 55*). Step 3: To define the hierarchical structure of the directed TF network, we used a graph-theoretical algorithm called *vertex-sort* (*33*), which identifies strongly connected components and applies the leaf removal algorithm on the graph and on its transpose which can identify the precise topological ordering of members in any directed network based on the number of connections that start from or end at each TF, indicating whether a TF is more regulating or more regulated. This allows for an approximate stratification of TFs within each dataset. Edges and nodes in the network were visualized by igraph R package (https://igraph.org/r/). Centrality statistics of each TF node was calculated using qgraph R package *centrality_auto ()* function.

### RNA-seq library preparation

RNA from FACS-sorted neurons of the sensorimotor cortex (∼1000 cells) was isolated with the NucleoSpin RNA XS kit (CloneTech) according to the manufacturer’s protocol. RNA-seq libraries for cortical motor neurons were prepared with the QuantSeq 3’mRNA-Seq library prep kit FWD for Illumina (Lexogen) following the manufacturer’s instructions, while RNA-seq libraries for RGCs were generated using TruSeq with RiboZero gold following the manufacturers’s instructions. The cDNA was fragmented to 300 base pairs (bp) using the Covaris M220 (Covaris), and then the manufacturer’s instructions were followed for end repair, adaptor ligation, and library amplification. The libraries were quantified by the Qubit dsDNA HS Assay Kit (Molecular Probes); Library size distribution and molar concentration of cDNA molecules in each library were determined by the Agilent High Sensitivity DNA Assay on an Agilent 2200 TapeStation system. Libraries were multiplexed into a single pool and sequenced using a HiSeq4000 instrument (Illumina, San Diego, CA) to generate 69 bp single-end reads. The average read depth for each library is ∼11 million for cortical motor neurons and ∼33 million for RGCs.

### RNA-seq read alignment and processing

Sensorimotor cortex neuronal RNA-seq data were mapped to the reference genome (mm10 / GRCm38) using STAR (*153*). Alignment and duplication metrics were collected using PICARD tools functions CollectRnaSeqMetrics and MarkDuplicates respectively (http://broadinstitute.github.io/picard/). Transcript abundance from aligned reads were quantified by Salmon (*154*), followed by summarization to the gene level using the R package Tximport (*155*). Sequencing depth was normalized between samples using geometric mean (GEO) in DESeq2 package (*156*). Removal of unwanted variation (RUV) was used to remove batch effects (*157*) and genes with no counts in over 50% of the samples were removed.

### Gene set enrichment analysis (GSEA)

GSEA v2.0 software with default settings (*98*) was used to identify upstream TFs of the genes associated with the combined pro-regenerative treatments of AAV2-sh.*pten*, Oncomodulin plus CPT-cAMP. These genes were ranked by their correlations of expression changes with treatments measured by directional p-value, which is calculated as -sign(log Treatment/Control)*(log10 p-value). A positive correlation indicates up-regulation of a gene by pro-regenerative treatment, while a negative correlation indicates down-regulation. A total of 1137 gene sets known to be targeted by transcription factors were downloaded from MsigDB (v5.1), and each set of the TF target genes were compared to the genes associated with the pro-regenerative treatments. An enrichment score (ES) is returned for each comparison, which represents the degree to which the TF-target list is over-represented at the top or bottom of the ranked gene list. The score is calculated by walking down the gene list, increasing a running-sum statistic when we encounter a gene in the TF-target list and decreasing when it is not. The magnitude of the increment depends on the gene statistics so as to determine whether a specific set of a TF’s target genes is randomly distributed throughout genes of interest, or primarily found at the top or bottom.

### Differential gene expression

Principle component analysis (PCA) of the normalized expression data (first five PCs) was correlated with potential technical covariates, including sex, aligning and sequencing bias calculated from STAR and Picard respectively. Differential gene expression by limma voom (*158*) was performed on normalized gene counts, including the first two PCs of aligning and sequencing bias as covariates: ∼ Genotype + AlignSeq.PC1 + AlignSeq.PC2. Differentially expressed genes were determined at FDR p value < 0.1 (Supplemental Table 1). Gene overlap analysis between DEGs and REST targeted gene sets was performed using the R package GeneOverlap. One-tailed P values were used (equivalent to hypergeometric P value) since we do not assume enrichment a priori.

### Gene Ontology Analysis

GO term enrichment analysis was performed using the gProfileR package (*159*) and Ingenuity Pathway Analysis (IPA) Software (Qiagen), using expressed genes in each of the normalized dataset as background. A maximum of top 10 canonical biological pathways, disease and function from each analysis were chosen from GO terms with FDR of p values < 0.05 and at least 10 genes overlapping the test data. The R package clusterProfiler (*160*) was used to plot the DEGs connecting to a specific GO term, with source code modified to accept GO terms from gProfileR and IPA.

### Weighted gene co-expression network analysis

Sequencing and aligning covariates were regressed out from normalized expression data using a linear model. Co-expression network was constructed using the WGCNA package (*82*). Briefly, pair-wise Pearson correlations between each gene pair were calculated and transformed to a signed adjacency matrix using a power of 10, as it was the smallest threshold that resulted in a scale-free R^2^ fit of 0.8. The adjacency matrix was used to construct a topological overlap dissimilarity matrix, from which hierarchical clustering of genes as modules were determined by a dynamic tree-cutting algorithm (Supplemental Table 3).

### WGCNA module annotation

To classify up- or down-regulated modules, the module eigengene, defined as the first principle component of a module that explains the maximum possible variability of that module, was related to genotype (wild-type vs REST cKO) using a linear model. Modules were considered to be significantly associated with the phenotype when Bonferroni corrected p values are less than 0.05. As a first step towards functional annotation, a hypergeometric analysis was used to examine each module’s association with the regeneration-associated gene (RAGs) module known to be activated by peripheral injury (Chandran et al., 2016). Modules were considered to be significantly associated with the RAG program when Bonferroni corrected p values are less than 0.05. To further annotate modules at a general level, we applied gene ontology (GO) enrichment analyses on each module. We also calculated Pearson correlations between each gene and each module eigengene as a gene’s module membership (Supplemental Table 3), and hub genes were defined as being those with highest correlations (kME > 0.7), which represent the most central genes in the co-expression network.

### Protein-protein interaction (PPI) network analysis

We established interactions of proteins encoded by genes from each of the co-expression modules (RESTUP1 [202 genes], RESTUP3 [636 genes], and RAG module [286 genes]) using InWeb database, which combines reported protein interactions from MINT, BIND, IntAct, KEGG annotated protein-protein interactions (PPrel), KEGG Enzymes involved in neighboring steps (ECrel), and Reactome (*161, 162*). The significance of PPIs within the network was further determined by DAPPLE, which uses a within-degree within-node permutation method that allows us to rank PPI hubs by P value. The PPI networks were visualized by igraph R package (https://igraph.org/r/), or Ingenuity Pathway Analysis (IPA) Software (Qiagen)

## Acknowledgments

We are grateful for the support of the Dr. Miriam and Sheldon G. Adelson Medical Research Foundation (DHG, LB, CJW, MS), the National Eye Institute (U01EY027261-01 to LB, JLG), U.S. Department of Defense (CDMRP W81XWH-16-1-0043 to LB, JLG), and NICHD IDDRC HD018655 (Imaging, Cell-Sorting, to CJW,LB). We also wish to thank Dr. Gail Mandel (Oregon Health Sciences Univ.), Drs. Mihaela Stavarache and Michael Kaplitt (Weill Cornell Medical College) for generously providing viral vectors and advice, and Dr. Jenny Hsieh (University of Texas at San Antonio) for providing the initial REST^flx/flx^ breeder mice.

## Author Contributions

Y.C., Y.Y., M.V.S., L.I.B. and D.H.G. designed and directed the experiments and guided the analysis. Y.C., Y.Y., L.I.B. and D.H.G. prepared the figures and wrote the manuscript. Y.C. performed bioinformatic analyses on the RNA-seq datasets of cortical motor neurons and RGCs. A.Z. performed transcription factor network analysis on the microarray datasets. Y.C. and A.Z. performed experiments on mouse DRG cultures with guidance from C.J.W. Y.Y. performed experiments on mouse RGC cultures and 293T cells. Y.C., A.M.B., K.G., Y.A., and K.P. performed SCI experiments and collected brain and spinal cord samples for immunostaining. Y.C., K.G. and J. Ou processed cortical motor neurons and prepared RNA-seq libraries. Y.Y. and H.Y.G. conducted optic nerve crush experiments, processed retina for immunostaining and RGCs for RT-PCR and RNA-seq. R.K. performed initial processing of RNA-seq data from RGCs. Y.C., A.Z., and C.J.K. bred the REST^flx/flx^ and REST^flx/flx^; STOP^flx/flx^TdTomato mice. All authors discussed the results and provided comments and revisions on the manuscript.

## Ethics Declarations

The authors declare no competing interests.

## Data Availability

The accession numbers for the data generated in this paper are GSE141583 and GSE142881.

## Code Availability

All code for processing and analyzing the data presented in this work are available upon request.

## Supplemental Tables

**Table S1**. Differentially expressed genes (DEGs) comparing wild-type and REST knockout cortical motor neurons at 0, 1, 3, 7 days after SCI.

**Table S2**. Annotation of molecules in the regeneration-associated protein-protein interaction network in Figure 4D.

**Table S3**. Module eigengenes (MEs) of co-expression gene networks and module membership of each gene in RNA-seq of wild-type or REST knockout cortical motor neurons in sham or SCI conditions.

**Table S4**. Expression level changes of REST-repressed genes predicted by ARACNe comparing RGCs sorted at 1, 3, 5 days after optic nerve crush with pro-regenerative treatment to non-regenerative RGCs with control treatment.

## References

1. S. R. Cajal, Cajal’s degeneration and regeneration of the nervous system. (History of Neuroscience, 1991).

2. Z. He, Y. Jin, Intrinsic Control of Axon Regeneration. Neuron 90, 437–451 (2016).

3. J. W. Fawcett, J. Verhaagen, Intrinsic Determinants of Axon Regeneration. Dev Neurobiol 78, 890–897 (2018).

4. J. Silver, J. H. Miller, Regeneration beyond the glial scar. Nature Reviews Neuroscience 5, 146–156 (2004).

5. M. T. Filbin, Myelin-associated inhibitors of axonal regeneration in the adult mammalian CNS. Nat Rev Neurosci 4, 703–713 (2003).

6. N. Klapka, H. W. Muller, Collagen matrix in spinal cord injury. J Neurotrauma 23, 422–435 (2006).

7. T. M. O’Shea, J. E. Burda, M. V. Sofroniew, Cell biology of spinal cord injury and repair. J Clin Invest 127, 3259–3270 (2017).

8. M. A. Anderson et al., Required growth facilitators propel axon regeneration across complete spinal cord injury. Nature 561, 396 (2018).

9. B. Zheng, et al., Genetic deletion of the Nogo receptor does not reduce neurite inhibition in vitro or promote corticospinal tract regeneration in vivo. (2005).

10. J. K. Lee et al., Combined genetic attenuation of myelin and semaphorin-mediated growth inhibition is insufficient to promote serotonergic axon regeneration. J Neurosci 30, 10899–10904 (2010).

11. D. Fischer, V. Petkova, S. Thanos, L. I. Benowitz, Switching mature retinal ganglion cells to a robust growth state in vivo: gene expression and synergy with RhoA inactivation. J. Neurosci. 24, 8726–8740 (2004).

12. D. Fischer, Z. He, L. I. Benowitz, Counteracting the Nogo receptor enhances optic nerve regeneration if retinal ganglion cells are in an active growth state. J Neurosci 24, 1646–1651 (2004).

13. T. L. Dickendesher et al., NgR1 and NgR3 are receptors for chondroitin sulfate proteoglycans. Nat Neurosci 15, 703–712 (2012).

14. K. K. Park et al., Promoting axon regeneration in the adult CNS by modulation of the PTEN/mTOR pathway. Science 322, 963–966 (2008).

15. T. Kurimoto et al., Long-distance axon regeneration in the mature optic nerve: contributions of oncomodulin, cAMP, and pten gene deletion. J Neurosci 30, 15654–15663 (2010).

16. S. de Lima et al., Full-length axon regeneration in the adult mouse optic nerve and partial recovery of simple visual behaviors. Proc Natl Acad Sci U S A 109, 9149–9154 (2012).

17. F. Sun et al., Sustained axon regeneration induced by co-deletion of PTEN and SOCS3. Nature, (2012).

18. N. Abe, V. Cavalli, Nerve injury signaling. Curr Opin Neurobiol 18, 276–283 (2008).

19. M. Mahar, V. Cavalli, Intrinsic mechanisms of neuronal axon regeneration. Nat Rev Neurosci 19, 323–337 (2018).

20. V. Chandran et al., A Systems-Level Analysis of the Peripheral Nerve Intrinsic Axonal Growth Program. Neuron 89, 956–970 (2016).

21. F. M. Bareyre et al., In vivo imaging reveals a phase-specific role of STAT3 during central and peripheral nervous system axon regeneration. Proc Natl Acad Sci U S A 108, 6282–6287 (2011).

22. D. L. Moore et al., KLF family members regulate intrinsic axon regeneration ability. Science 326, 298–301 (2009).

23. M. G. Blackmore et al., Kruppel-like Factor 7 engineered for transcriptional activation promotes axon regeneration in the adult corticospinal tract. Proc Natl Acad Sci U S A 109, 7517–7522 (2012).

24. Z. Wang, A. Reynolds, A. Kirry, C. Nienhaus, M. G. Blackmore, Overexpression of Sox11 promotes corticospinal tract regeneration after spinal injury while interfering with functional recovery. J Neurosci 35, 3139–3145 (2015).

25. M. W. Norsworthy et al., Sox11 Expression Promotes Regeneration of Some Retinal Ganglion Cell Types but Kills Others. Neuron 94, 1112–1120.e1114 (2017).

26. M. M. Babu, N. M. Luscombe, L. Aravind, M. Gerstein, S. A. Teichmann, Structure and evolution of transcriptional regulatory networks. Curr Opin Struct Biol 14, 283–291 (2004).

27. A. Blais, B. D. Dynlacht, Constructing transcriptional regulatory networks. Genes Dev 19, 1499–1511 (2005).

28. L. Ni et al., Dynamic and complex transcription factor binding during an inducible response in yeast. Genes Dev 23, 1351–1363 (2009).

29. G. May et al., Dynamic analysis of gene expression and genome-wide transcription factor binding during lineage specification of multipotent progenitors. Cell Stem Cell 13, 754–768 (2013).

30. M. B. Gerstein et al., Architecture of the human regulatory network derived from ENCODE data. Nature 489, 91 (2012).

31. N. D. Fagoe, J. van Heest, J. Verhaagen, Spinal cord injury and the neuron-intrinsic regeneration-associated gene program. Neuromolecular medicine 16, 799–813 (2014).

32. J. Kim, J. Chu, X. Shen, J. Wang, S. H. Orkin, An extended transcriptional network for pluripotency of embryonic stem cells. Cell 132, 1049–1061 (2008).

33. R. Jothi et al., Genomic analysis reveals a tight link between transcription factor dynamics and regulatory network architecture. Mol Syst Biol 5, 294 (2009).

34. A. P. Boyle et al., Comparative analysis of regulatory information and circuits across distant species. Nature 512, 453–456 (2014).

35. H. Yu, M. Gerstein, Genomic analysis of the hierarchical structure of regulatory networks. (2006).

36. A. Blesch et al., Conditioning lesions before or after spinal cord injury recruit broad genetic mechanisms that sustain axonal regeneration: superiority to camp-mediated effects. Experimental neurology 235, 162–173 (2012).

37. R. S. Griffin et al., Complement induction in spinal cord microglia results in anaphylatoxin C5a-mediated pain hypersensitivity. J Neurosci 27, 8699–8708 (2007).

38. B. Yu et al., miR-182 inhibits Schwann cell proliferation and migration by targeting FGF9 and NTM, respectively at an early stage following sciatic nerve injury. Nucleic Acids Res 40, 10356–10365 (2012).

39. I. Michaelevski et al., Signaling to transcription networks in the neuronal retrograde injury response. Sci Signal 3, ra53 (2010).

40. J. Ryge et al., in BMC Genomics. (2010), vol. 11, pp. 365.

41. A. De Biase et al., Gene expression profiling of experimental traumatic spinal cord injury as a function of distance from impact site and injury severity. Physiol Genomics 22, 368–381 (2005).

42. J. A. Chong et al., REST: a mammalian silencer protein that restricts sodium channel gene expression to neurons. Cell 80, 949–957 (1995).

43. C. J. Schoenherr, D. J. Anderson, The neuron-restrictive silencer factor (NRSF): a coordinate repressor of multiple neuron-specific genes. Science 267, 1360–1363 (1995).

44. N. Ballas, C. Grunseich, D. D. Lu, J. C. Speh, G. Mandel, REST and its corepressors mediate plasticity of neuronal gene chromatin throughout neurogenesis. Cell 121, 645–657 (2005).

45. N. Ballas, G. Mandel, The many faces of REST oversee epigenetic programming of neuronal genes. Curr Opin Neurobiol 15, 500–506 (2005).

46. T. Nechiporuk et al., The REST remodeling complex protects genomic integrity during embryonic neurogenesis. Elife 5, e09584 (2016).

47. J. C. McGann et al., Polycomb- and REST-associated histone deacetylases are independent pathways toward a mature neuronal phenotype. Elife 3, e04235 (2014).

48. A. A. Margolin et al., in BMC Bioinformatics. (2006), vol. 7, pp. S7.

49. A. Lachmann, F. M. Giorgi, G. Lopez, A. Califano, ARACNe-AP: gene network reverse engineering through adaptive partitioning inference of mutual information. Bioinformatics 32, 2233–2235 (2016).

50. X. Zhao et al., The N-Myc-DLL3 cascade is suppressed by the ubiquitin ligase Huwe1 to inhibit proliferation and promote neurogenesis in the developing brain. Dev Cell 17, 210–221 (2009).

51. M. S. Carro et al., The transcriptional network for mesenchymal transformation of brain tumours. Nature 463, 318–325 (2010).

52. C. Lefebvre et al., A human B-cell interactome identifies MYB and FOXM1 as master regulators of proliferation in germinal centers. Mol Syst Biol 6, 377 (2010).

53. G. Della Gatta et al., Reverse engineering of TLX oncogenic transcriptional networks identifies RUNX1 as tumor suppressor in T-ALL. Nature medicine 18, 436–440 (2012).

54. R. Kushwaha et al., Interrogation of a Context-Specific Transcription Factor Network Identifies Novel Regulators of Pluripotency. Stem Cells 33, 367–377 (2015).

55. A. Lachmann et al., ChEA: transcription factor regulation inferred from integrating genome-wide ChIP-X experiments. Bioinformatics 26, 2438–2444 (2010).

56. S. G. Landt et al., ChIP-seq guidelines and practices of the ENCODE and modENCODE consortia. Genome Res 22, 1813–1831 (2012).

57. H. Tsujino et al., Activating transcription factor 3 (ATF3) induction by axotomy in sensory and motoneurons: A novel neuronal marker of nerve injury. Mol Cell Neurosci 15, 170–182 (2000).

58. R. Seijffers, C. D. Mills, C. J. Woolf, in J Neurosci. (2007), vol. 27, pp. 7911–7920.

59. R. Jenkins, S. P. Hunt, Long-term increase in the levels of c-jun mRNA and jun protein-like immunoreactivity in motor and sensory neurons following axon damage. Neurosci Lett 129, 107–110 (1991).

60. F. W. Schwaiger et al., Peripheral but not central axotomy induces changes in Janus kinases (JAK) and signal transducers and activators of transcription (STAT). Eur J Neurosci 12, 1165–1176 (2000).

61. G. Raivich et al., The AP-1 transcription factor c-Jun is required for efficient axonal regeneration. Neuron 43, 57–67 (2004).

62. H. Zou, C. Ho, K. Wong, M. Tessier-Lavigne, Axotomy-induced Smad1 activation promotes axonal growth in adult sensory neurons. J Neurosci 29, 7116–7123 (2009).

63. Saijilafu et al., PI3K-GSK3 signalling regulates mammalian axon regeneration by inducing the expression of Smad1. Nat Commun 4, 2690 (2013).

64. M. P. Jankowski, P. K. Cornuet, S. McIlwrath, H. R. Koerber, K. M. Albers, SRY-box containing gene 11 (Sox11) transcription factor is required for neuron survival and neurite growth. Neuroscience 143, 501–514 (2006).

65. X. Jing, T. Wang, S. Huang, J. C. Glorioso, K. M. Albers, The transcription factor Sox11 promotes nerve regeneration through activation of the regeneration-associated gene Sprr1a. Experimental neurology 233, 221–232 (2012).

66. W. Renthal et al., Transcriptional Reprogramming of Distinct Peripheral Sensory Neuron Subtypes after Axonal Injury. Neuron, (2020).

67. R. Deumens, G. C. Koopmans, E. A. Joosten, Regeneration of descending axon tracts after spinal cord injury. Prog Neurobiol 77, 57–89 (2005).

68. M. V. Sofroniew, Dissecting spinal cord regeneration. Nature 557, 343 (2018).

69. D. H. Geschwind, G. Konopka, Neuroscience in the era of functional genomics and systems biology. Nature 461, 908–915 (2009).

70. A. Tedeschi, Tuning the Orchestra: Transcriptional Pathways Controlling Axon Regeneration. Frontiers in molecular neuroscience 4, (2011).

71. L. Aigner et al., Overexpression of the neural growth-associated protein GAP-43 induces nerve sprouting in the adult nervous system of transgenic mice. Cell 83, 269–278 (1995).

72. H. M. Bomze, K. R. Bulsara, B. J. Iskandar, P. Caroni, J. H. Skene, Spinal axon regeneration evoked by replacing two growth cone proteins in adult neurons. Nature neuroscience 4, 38–43 (2001).

73. G. Keilhoff, S. Wiegand, H. Fansa, Vav deficiency impedes peripheral nerve regeneration in mice. Restor Neurol Neurosci 30, 463–479 (2012).

74. L. F. Gumy et al., Transcriptome analysis of embryonic and adult sensory axons reveals changes in mRNA repertoire localization. RNA 17, 85–98 (2011).

75. N. Kishimoto, K. Shimizu, K. Sawamoto, in Dis Model Mech. (2012), vol. 5, pp. 200–209.

76. A. Guijarro-Belmar et al., Epac2 Elevation Reverses Inhibition by Chondroitin Sulfate Proteoglycans In Vitro and Transforms Postlesion Inhibitory Environment to Promote Axonal Outgrowth in an Ex Vivo Model of Spinal Cord Injury. J Neurosci 39, 8330–8346 (2019).

77. N. Kato et al., in Journal of neuroinflammation. (2013), vol. 10, pp. 1.

78. A. W. Bruce et al., Genome-wide analysis of repressor element 1 silencing transcription factor/neuron-restrictive silencing factor (REST/NRSF) target genes. Proc Natl Acad Sci U S A 101, 10458–10463 (2004).

79. R. Johnson et al., in PLoS Biol. (2008), vol. 6.

80. V. Matys et al., TRANSFAC: transcriptional regulation, from patterns to profiles. Nucleic Acids Res 31, 374–378 (2003).

81. S. Mukherjee, R. Brulet, L. Zhang, J. Hsieh, REST regulation of gene networks in adult neural stem cells. Nat Commun 7, 13360 (2016).

82. P. Langfelder, S. Horvath, in BMC Bioinformatics. (2008), vol. 9, pp. 559.

83. B. Zhang, S. Horvath, A general framework for weighted gene co-expression network analysis. Stat Appl Genet Mol Biol 4, Article17 (2005).

84. A. J. Aguayo et al., Degenerative and regenerative responses of injured neurons in the central nervous system of adult mammals. Philos Trans R Soc Lond B Biol Sci 331, 337–343 (1991).

85. T. Kurimoto et al., Neutrophils express oncomodulin and promote optic nerve regeneration. J Neurosci 33, 14816–14824 (2013).

86. Y. Yin et al., Oncomodulin links inflammation to optic nerve regeneration. Proc Natl Acad Sci U S A 106, 19587–19592 (2009).

87. Y. Yin et al., Oncomodulin is a macrophage-derived signal for axon regeneration in retinal ganglion cells. Nat Neurosci 9, 843–852 (2006).

88. Y. Yin, et al., Stromal cell-derived factor-1 (SDF-1) contributes to inflammation-induced optic nerve regeneration and retinal ganglion cell survival. Program No. 531.04. 2012 Neuroscience Meeting Planner. New Orleans, LA: Society for Neuroscience, 2012, Online., (2012).

89. V. Pernet et al., Long-distance axonal regeneration induced by CNTF gene transfer is impaired by axonal misguidance in the injured adult optic nerve. Neurobiology of disease 51, 202–213 (2013).

90. A. Apara et al., KLF9 and JNK3 Interact to Suppress Axon Regeneration in the Adult CNS. J Neurosci 37, 9632–9644 (2017).

91. E. F. Trakhtenberg et al., Zinc chelation and Klf9 knockdown cooperatively promote axon regeneration after optic nerve injury. Exp Neurol 300, 22–29 (2018).

92. Y. Li et al., Mobile zinc increases rapidly in the retina after optic nerve injury and regulates ganglion cell survival and optic nerve regeneration. Proc Natl Acad Sci U S A 114, E209–E218 (2017).

93. J. H. Lim et al., Neural activity promotes long-distance, target-specific regeneration of adult retinal axons. Nat Neurosci 19, 1073–1084 (2016).

94. L. I. Benowitz, Z. He, J. L. Goldberg, Reaching the brain: Advances in optic nerve regeneration. Experimental neurology 287, 365–373 (2017).

95. Y. Zhang et al., Elevating Growth Factor Responsiveness and Axon Regeneration by Modulating Presynaptic Inputs. Neuron, (2019).

96. G. Feng et al., Imaging neuronal subsets in transgenic mice expressing multiple spectral variants of GFP. Neuron 28, 41–51 (2000).

97. M. A. Stavarache, S. Musatov, M. McGill, M. Vernov, M. G. Kaplitt, The tumor suppressor PTEN regulates motor responses to striatal dopamine in normal and Parkinsonian animals. Neurobiology of disease 82, 487–494 (2015).

98. A. Subramanian et al., Gene set enrichment analysis: a knowledge-based approach for interpreting genome-wide expression profiles. Proc Natl Acad Sci U S A 102, 15545–15550 (2005).

99. I. Yevshin, R. Sharipov, S. Kolmykov, Y. Kondrakhin, F. Kolpakov, in Nucleic Acids Res. (2019), vol. 47, pp. D100–105.

100. J. F. Borisoff et al., Suppression of Rho-kinase activity promotes axonal growth on inhibitory CNS substrates. Mol Cell Neurosci 22, 405–416 (2003).

101. Z. L. Chen, S. Strickland, Laminin gamma1 is critical for Schwann cell differentiation, axon myelination, and regeneration in the peripheral nerve. The Journal of cell biology 163, 889–899 (2003).

102. C. L. Tan, J. C. F. Kwok, R. Patani, S. Chandran, J. W. Fawcett, Integrin activation promotes axon growth on inhibitory CSPGs by enhancing integrin signaling. J Neurosci 31, 6289–6295 (2011).

103. B. Lang et al., Modulation of the proteoglycan receptor PTPσ promotes recovery after spinal cord injury. Nature 518, 404–408 (2015).

104. V. Lisi et al., Enhanced Neuronal Regeneration in the CAST/Ei Mouse Strain Is Linked to Expression of Differentiation Markers after Injury. Cell Rep 20, 1136–1147 (2017).

105. D. S. Smith, J. H. Skene, A transcription-dependent switch controls competence of adult neurons for distinct modes of axon growth. J Neurosci 17, 646–658 (1997).

106. Saijilafu, F. Q. Zhou, Genetic study of axon regeneration with cultured adult dorsal root ganglion neurons. J Vis Exp, (2012).

107. Z. Gao et al., in J Neurosci. (2011), vol. 31, pp. 9772–9786.

108. K. Liu et al., PTEN deletion enhances the regenerative ability of adult corticospinal neurons. Nature neuroscience 13, 1075–1081 (2010).

109. M. A. Anderson et al., Astrocyte scar formation aids central nervous system axon regeneration. Nature 532, 195 (2016).

110. C. G. Geoffroy, B. Zheng, Myelin-Associated Inhibitors in Axonal Growth After CNS Injury. Curr Opin Neurobiol 0, 31–38 (2014).

111. G. Mandel et al., Repressor element 1 silencing transcription factor (REST) controls radial migration and temporal neuronal specification during neocortical development. Proc Natl Acad Sci U S A 108, 16789–16794 (2011).

112. Y. Yin et al., Macrophage-derived factors stimulate optic nerve regeneration. Journal of Neuroscience 23, 2284–2293 (2003).

113. M. P. Jankowski et al., Sox11 transcription factor modulates peripheral nerve regeneration in adult mice. Brain Res 1256, 43–54 (2009).

114. S. T. Mehta, X. Luo, K. K. Park, J. L. Bixby, V. P. Lemmon, Hyperactivated Stat3 boosts axon regeneration in the CNS. Experimental neurology 280, 115–120 (2016).

115. C. Lindwall, M. Kanje, Retrograde axonal transport of JNK signaling molecules influence injury induced nuclear changes in p-c-Jun and ATF3 in adult rat sensory neurons. Mol Cell Neurosci 29, 269–282 (2005).

116. J. Qiu, W. B. Cafferty, S. B. McMahon, S. W. Thompson, Conditioning injury-induced spinal axon regeneration requires signal transducer and activator of transcription 3 activation. J Neurosci 25, 1645–1653 (2005).

117. P. D. Smith et al., SOCS3 deletion promotes optic nerve regeneration in vivo. Neuron 64, 617–623 (2009).

118. C. J. Schoenherr, A. J. Paquette, D. J. Anderson, Identification of potential target genes for the neuron-restrictive silencer factor. Proc Natl Acad Sci U S A 93, 9881–9886 (1996).

119. C. Conaco, S. Otto, J. J. Han, G. Mandel, Reciprocal actions of REST and a microRNA promote neuronal identity. Proc Natl Acad Sci U S A 103, 2422–2427 (2006).

120. S. J. Otto et al., A new binding motif for the transcriptional repressor REST uncovers large gene networks devoted to neuronal functions. J Neurosci 27, 6729–6739 (2007).

121. A. Calderone et al., Ischemic insults derepress the gene silencer REST in neurons destined to die. J Neurosci 23, 2112–2121 (2003).

122. K. M. Noh et al., Repressor element-1 silencing transcription factor (REST)-dependent epigenetic remodeling is critical to ischemia-induced neuronal death. Proc Natl Acad Sci U S A 109, E962–971 (2012).

123. K. Palm, N. Belluardo, M. Metsis, T. Timmusk, Neuronal expression of zinc finger transcription factor REST/NRSF/XBR gene. J Neurosci 18, 1280–1296 (1998).

124. M. Garriga-Canut et al., 2-Deoxy-D-glucose reduces epilepsy progression by NRSF-CtBP-dependent metabolic regulation of chromatin structure. Nature neuroscience 9, 1382–1387 (2006).

125. S. McClelland et al., Neuron-restrictive silencer factor-mediated hyperpolarization-activated cyclic nucleotide gated channelopathy in experimental temporal lobe epilepsy. Ann Neurol 70, 454–464 (2011).

126. S. McClelland et al., The transcription factor NRSF contributes to epileptogenesis by selective repression of a subset of target genes. Elife 3, e01267 (2014).

127. C. Zuccato et al., Huntingtin interacts with REST/NRSF to modulate the transcription of NRSE-controlled neuronal genes. Nat Genet 35, 76–83 (2003).

128. T. Lu et al., REST and stress resistance in ageing and Alzheimer’s disease. Nature 507, 448–454 (2014).

129. J. M. Zullo et al., Regulation of lifespan by neural excitation and REST. Nature 574, 359–364 (2019).

130. H. Uchida, L. Ma, H. Ueda, Epigenetic gene silencing underlies C-fiber dysfunctions in neuropathic pain. J Neurosci 30, 4806–4814 (2010).

131. D. E. Willis, M. Wang, E. Brown, L. Fones, J. W. Cave, Selective repression of gene expression in neuropathic pain by the neuron-restrictive silencing factor/repressor element-1 silencing transcription (NRSF/REST). Neurosci Lett 625, 20–25 (2016).

132. J. Zhang, S. R. Chen, H. Chen, H. L. Pan, RE1-silencing transcription factor controls the acute-to-chronic neuropathic pain transition and Chrm2 receptor gene expression in primary sensory neurons. J Biol Chem 293, 19078–19091 (2018).

133. Y. M. Oh, et al., Epigenetic regulator UHRF1 inactivates REST and growth suppressor gene expression via DNA methylation to promote axon regeneration. (2018).

134. M. H. Tuszynski, O. Steward, Concepts and Methods for the Study of Axonal Regeneration in the CNS. Neuron 74, 777–791 (2012).

135. M. B. Bunge, Bridging areas of injury in the spinal cord. Neuroscientist 7, 325–339 (2001).

136. H. Cheng, Y. Cao, L. Olson, Spinal cord repair in adult paraplegic rats: partial restoration of hind limb function. Science 273, 510–513 (1996).

137. P. Lu et al., Long-distance growth and connectivity of neural stem cells after severe spinal cord injury. Cell 150, 1264–1273 (2012).

138. K. Kadoya et al., Spinal cord reconstitution with homologous neural grafts enables robust corticospinal regeneration. Nature medicine 22, 479–487 (2016).

139. e. a. Liu K PTEN deletion enhances the regenerative ability of adult corticospinal neurons. - PubMed - NCBI. (2017).

140. J. C. Koch, E. Barski, P. Lingor, M. Bahr, U. Michel, Plasmids containing NRSE/RE1 sites enhance neurite outgrowth of retinal ganglion cells via sequestration of REST independent of NRSE dsRNA expression. FEBS J 278, 3472–3483 (2011).

141. C. Kole et al., Activating Transcription Factor 3 (ATF3) Protects Retinal Ganglion Cells and Promotes Functional Preservation After Optic Nerve Crush. Invest Ophthalmol Vis Sci 61, 31 (2020).

142. A. Brunet, S. R. Datta, M. E. Greenberg, Transcription-dependent and -independent control of neuronal survival by the PI3K-Akt signaling pathway. Curr Opin Neurobiol 11, 297–305 (2001).

143. T. Gu et al., CREB is a novel nuclear target of PTEN phosphatase. Cancer Res 71, 2821–2825 (2011).

144. K. Du, M. Montminy, CREB is a regulatory target for the protein kinase Akt/PKB. J Biol Chem 273, 32377–32379 (1998).

145. J. Satoh, N. Kawana, Y. Yamamoto, in Bioinform Biol Insights. (2013), vol. 7, pp. 357–368.

146. Y. Nakano et al., Defects in the Alternative Splicing-Dependent Regulation of REST Cause Deafness. Cell 174, 536–548.e521 (2018).

147. T. F. Westbrook et al., SCFbeta-TRCP controls oncogenic transformation and neural differentiation through REST degradation. Nature 452, 370–374 (2008).

148. M. Faronato et al., The deubiquitylase USP15 stabilizes newly synthesized REST and rescues its expression at mitotic exit. Cell Cycle 12, 1964–1977 (2013).

149. K. Zukor et al., Short hairpin RNA against PTEN enhances regenerative growth of corticospinal tract axons after spinal cord injury. J Neurosci 33, 15350–15361 (2013).

150. S. Leon, Y. Yin, J. Nguyen, N. Irwin, L. I. Benowitz, Lens injury stimulates axon regeneration in the mature rat optic nerve. J Neurosci 20, 4615–4626 (2000).

151. V. Swarup et al., Identification of evolutionarily conserved gene networks mediating neurodegenerative dementia. Nature medicine 25, 152–164 (2019).

152. M. A. Castro et al., Regulators of genetic risk of breast cancer identified by integrative network analysis. Nat Genet 48, 12–21 (2016).

153. A. Dobin et al., STAR: ultrafast universal RNA-seq aligner. Bioinformatics 29, 15–21 (2013).

154. R. Patro, G. Duggal, M. I. Love, R. A. Irizarry, C. Kingsford, Salmon: fast and bias-aware quantification of transcript expression using dual-phase inference. Nature methods 14, 417–419 (2017).

155. C. Soneson, M. I. Love, M. D. Robinson, Differential analyses for RNA-seq: transcript-level estimates improve gene-level inferences. F1000Res 4, 1521 (2015).

156. M. I. Love, W. Huber, S. Anders, in Genome Biol. (2014), vol. 15.

157. D. Risso, J. Ngai, T. P. Speed, S. Dudoit, Normalization of RNA-seq data using factor analysis of control genes or samples. Nature biotechnology 32, 896–902 (2014).

158. M. E. Ritchie et al., limma powers differential expression analyses for RNA-sequencing and microarray studies. Nucleic Acids Res 43, e47 (2015).

159. J. Reimand et al., g:Profiler—a web server for functional interpretation of gene lists (2016 update). Nucleic Acids Res 44, W83–89 (2016).

160. G. Yu, L. G. Wang, Y. Han, Q. Y. He, clusterProfiler: an R package for comparing biological themes among gene clusters. OMICS 16, 284–287 (2012).

161. A. Lundby et al., Annotation of loci from genome-wide association studies using tissue-specific quantitative interaction proteomics. Nature methods 11, 868–874 (2014).

162. E. J. Rossin et al., in PLoS Genet. (2011), vol. 7.

