## Supplemental materials for "Transcription factor network analysis identifies REST/NRSF as an intrinsic regulator of CNS regeneration"

### Figure S1

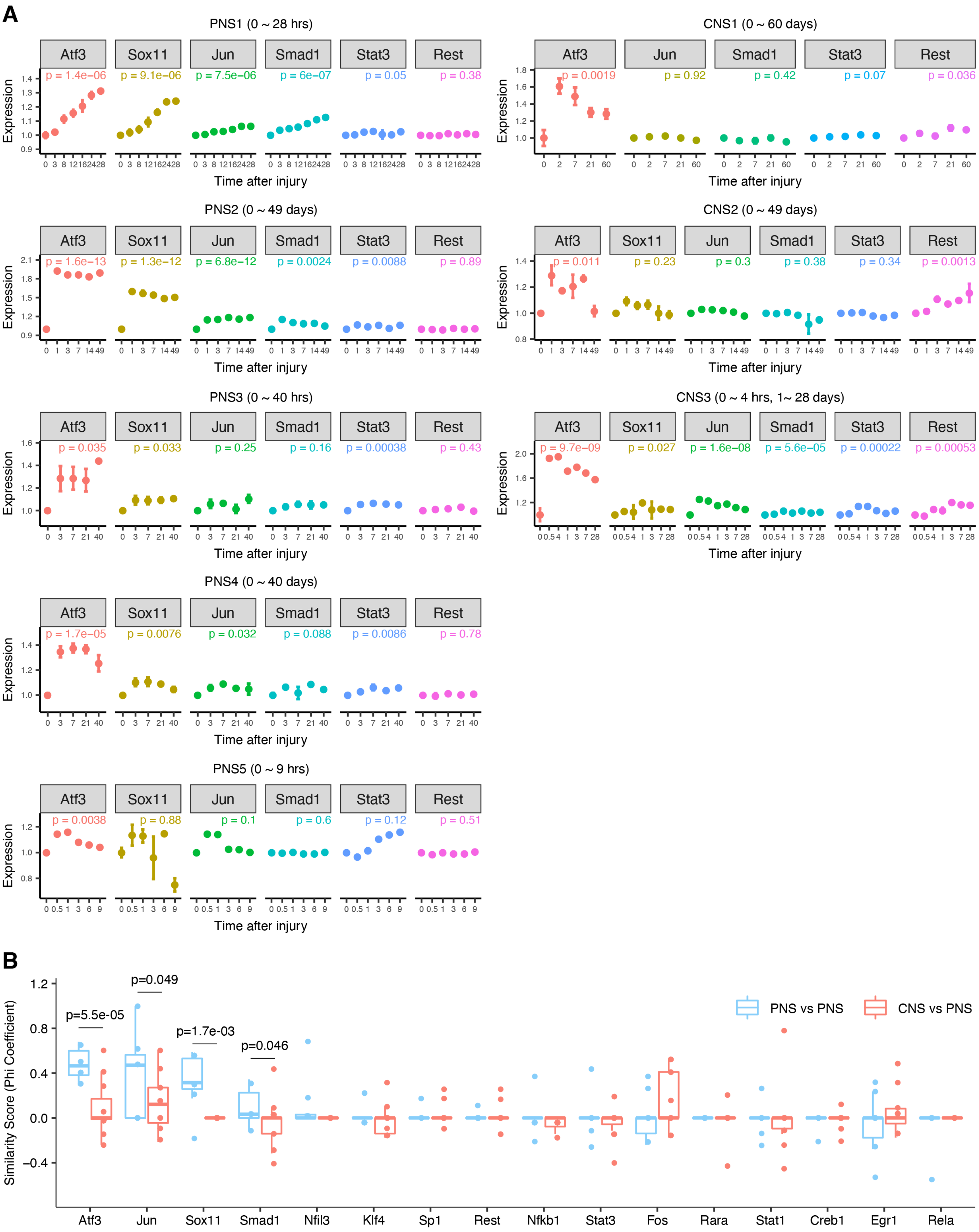
A
PNS1 (0 ~ 28 hrs)
Atf3
Sox11
Jun
Smad1
Stat3
Rest
p = 1.4e−06
p = 9.1e−06
p = 7.5e−06
p = 6e−07
p = 0.05
p = 0.38
Expression
Time after injury
CNS1 (0 ~ 60 days)
Atf3
Jun
Smad1
Stat3
Rest
p = 0.0019
p = 0.92
p = 0.42
p = 0.07
p = 0.036
Expression
Time after injury
PNS2 (0 ~ 49 days)
Atf3
Sox11
Jun
Smad1
Stat3
Rest
p = 1.6e−13
p = 1.3e−12
p = 6.8e−12
p = 0.0024
p = 0.0088
p = 0.89
Expression
Time after injury
CNS2 (0 ~ 49 days)
Atf3
Sox11
Jun
Smad1
Stat3
Rest
p = 0.011
p = 0.23
p = 0.3
p = 0.38
p = 0.34
p = 0.0013
Expression
Time after injury
PNS3 (0 ~ 40 hrs)
Atf3
Sox11
Jun
Smad1
Stat3
Rest
p = 0.035
p = 0.033
p = 0.25
p = 0.16
p = 0.00038
p = 0.43
Expression
Time after injury
CNS3 (0 ~ 4 hrs, 1~ 28 days)
Atf3
Sox11
Jun
Smad1
Stat3
Rest
p = 9.7e−09
p = 0.027
p = 1.6e−08
p = 5.6e−05
p = 0.00022
p = 0.00053
Expression
Time after injury
PNS4 (0 ~ 40 days)
Atf3
Sox11
Jun
Smad1
Stat3
Rest
p = 1.7e−05
p = 0.0076
p = 0.032
p = 0.088
p = 0.0086
p = 0.78
Expression
Time after injury
PNS5 (0 ~ 9 hrs)
Atf3
Sox11
Jun
Smad1
Stat3
Rest
p = 0.0038
p = 0.88
p = 0.1
p = 0.6
p = 0.12
p = 0.51
Expression
Time after injury
B
Similarity Score (Phi Coefficient)
p=5.5e-05
p=0.049
p=1.7e-03
p=0.046
PNS vs PNS
CNS vs PNS
Atf3
Jun
Sox11
Smad1
Nfil3
Klf4
Sp1
Rest
Nfkb1
Stat3
Fos
Rara
Stat1
Creb1
Egr1
Rela

**Figure S1. TF networks comparing PNS and CNS, related to Figure 2.** **(A)** Relative expression levels of key regenerative TFs (*Atf3, Sox11, Jun, Smad1, Stat3, Rest*) across multiple PNS and CNS injury datasets listed in Figure 2B. Each dot represents the mean log fold changes normalized to control sham condition. N = 3 in each time point and error bars indicates SEM. p-values are from ANOVA comparing injured with sham control samples, adjusted with TukeyHSD. **(B)** Similarity of ARACNe-identified TF networks across multiple PNS and CNS injury datasets listed in Figure 2B. Phi coefficient was used to calculate the correlation of each TF's networks between two datasets since a TF's connections with the other TFs are binary (1 means a connection exists between two TFs and 0 means no connection). Each dot represents the Phi coefficient of a TF network between two datasets. Blue dots indicate similarity of TF networks comparing PNS vs PNS, and red dots represent similarity comparing CNS vs PNS. Statistical differences were calculated by Welch two sample t-test on Fisher-transformed z values from Phi coefficient, as coefficient correlations are not normally distributed. Comparisons with no statistical difference ($p > 0.05$) were not labeled with its p-values.

#### Figure S2

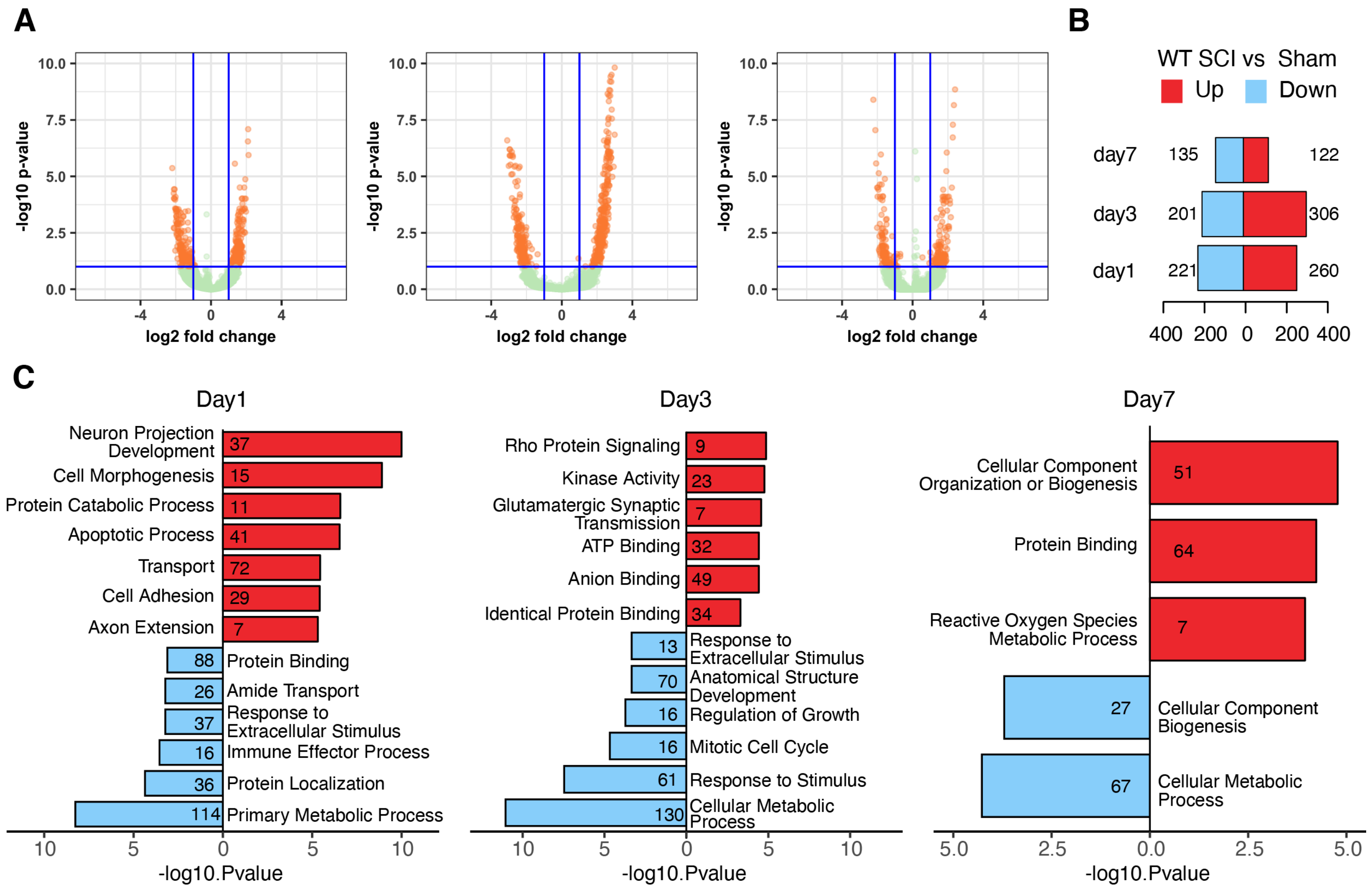
A
-log10 p-value
log2 fold change
B
WT SCI vs Sham
Up
Down
day7 135 122
day3 201 306
day1 221 260
400 200 0 200 400
C
Day1
Neuron Projection Development 37
Cell Morphogenesis 15
Protein Catabolic Process 11
Apoptotic Process 41
Transport 72
Cell Adhesion 29
Axon Extension 7
88 Protein Binding
26 Amide Transport
37 Response to Extracellular Stimulus
16 Immune Effector Process
36 Protein Localization
114 Primary Metabolic Process
-log10.Pvalue
Day3
Rho Protein Signaling 9
Kinase Activity 23
Glutamatergic Synaptic Transmission 7
ATP Binding 32
Anion Binding 49
Identical Protein Binding 34
13 Response to Extracellular Stimulus
70 Anatomical Structure Development
16 Regulation of Growth
16 Mitotic Cell Cycle
61 Response to Stimulus
130 Cellular Metabolic Process
-log10.Pvalue
Day7
Cellular Component Organization or Biogenesis 51
Protein Binding 64
Reactive Oxygen Species Metabolic Process 7
27 Cellular Component Biogenesis
67 Cellular Metabolic Process
-log10.Pvalue

**Figure S2. Differential gene expression analysis on RNA-seq of cortical neurons at 1, 3 and 7 days following SCI, related to Figure 3. (A)** Volcano plots showing differentially expressed genes (DEGs) at FDR p value < 0.1 at 1, 3, 7 and days after SCI compared to sham-treated group. **(B)** Number of DEGs at each time point. Up-regulated: red; Down-regulated genes: blue; |log2 FC| > 0.3). **(C)** Top gene ontology (GO) terms associated with DEGs at indicated condition (FDR adjusted enrichment p-value). A full list of DEGs are listed in Supplemental Table S1.

**Figure S3**

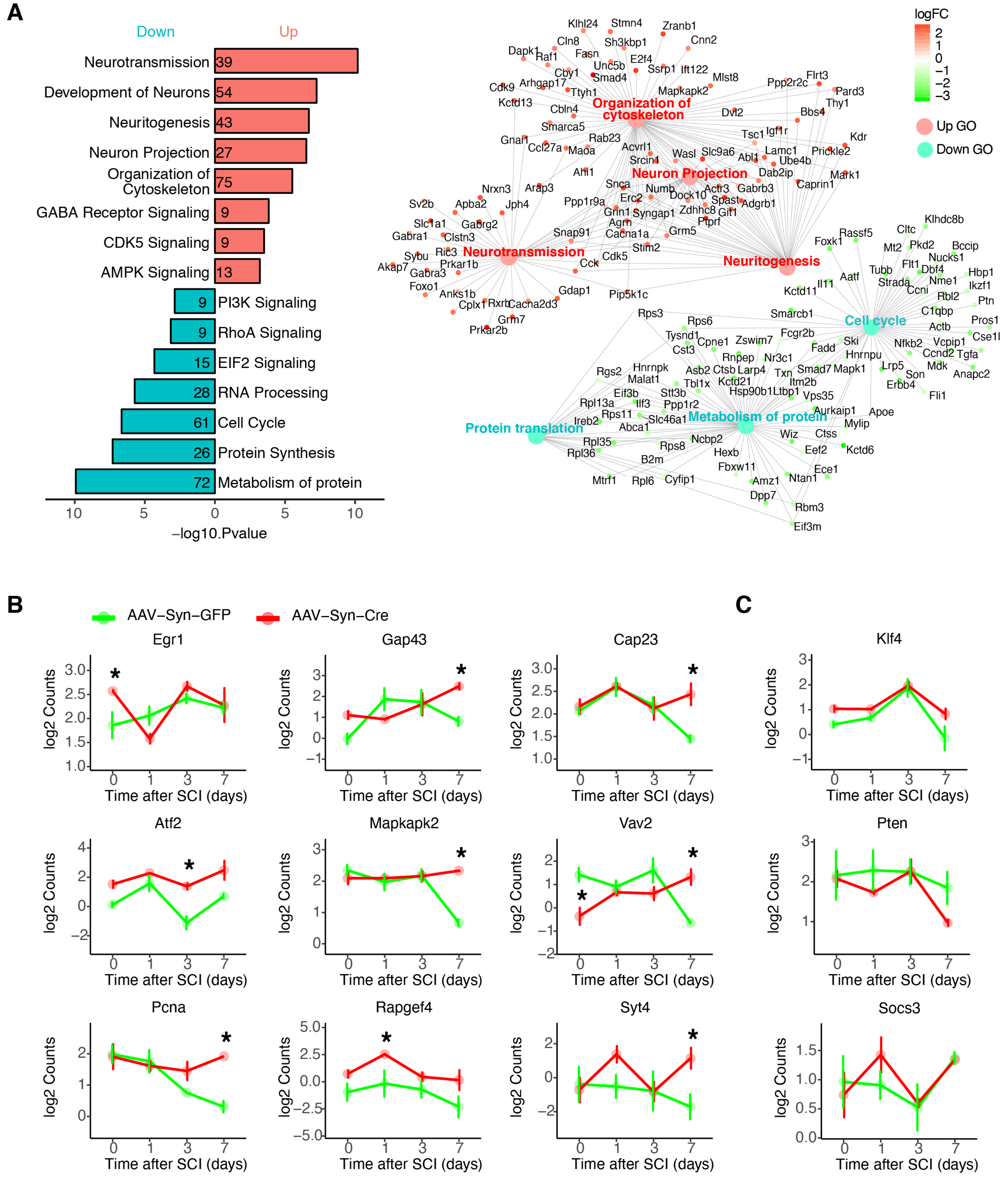
A
Down
Up
Neurotransmission 39
Development of Neurons 54
Neuritogenesis 43
Neuron Projection 27
Organization of Cytoskeleton 75
GABA Receptor Signaling 9
CDK5 Signaling 9
AMPK Signaling 13
9 PI3K Signaling
9 RhoA Signaling
15 EIF2 Signaling
28 RNA Processing
61 Cell Cycle
26 Protein Synthesis
72 Metabolism of protein
10 5 0 5 10
−log10.Pvalue
Organization of cytoskeleton
Neuron Projection
Neurotransmission
Neuritogenesis
Cell cycle
Protein translation
Metabolism of protein
logFC
2
1
0
−1
−2
−3
Up GO
Down GO
Klhl24 Stmn4 Zranb1 Cln8 Sh3kbp1 Cnn2 Dapk1 Raf1 Fasn E2f4 Unc5b Cby1 Ssrp1 Ift122 Smad4 Mlst8 Arhgap17 Cdk9 Ttyh1 Mapkapk2 Ppp2r2c Flrt3 Pard3 Thy1 Kctd13 Cbln4 Dvl2 Bbs4 Smarca5 Igf1r Tsc1 Kdr Gnai1 Rab23 Ccl27a Maoa Acvrl1 Lamc1 Prickle2 Srcin1 Wasl Slc9a6 Abl1 Ube4b Ahi1 Dab2ip Mark1 Caprin1 Nrxn3 Arap3 Snca Numb Actr3 Gabrb3 Dock10 Sv2b Apba2 Jph4 Ppp1r9a Erc2 Spast Adgrb1 Grin1 Syngap1 Zdhhc8 Git1 Slc1a1 Gabrg2 Agrn Ptprf Gabra1 Clstn3 Snap91 Cacna1a Grm5 Sybu Ric3 Stim2 Akap7 Prkar1b Cdk5 Gabra3 Cck Foxo1 Gdap1 Pip5k1c Anks1b Cplx1 Rxrb Cacna2d3 Grm7 Prkar2b
Klhdc8b Rassf5 Cltc Foxk1 Mt2 Pkd2 Bccip Nucks1 Flt1 Dbf4 Tubb Hbp1 Aatf Strada Nme1 Ikzf1 Kctd11 Il11 Ccni Rbl2 Ptn Smarcb1 C1qbp Pros1 Actb Cse1l Rps3 Rps6 Fcgr2b Nfkb2 Vcpip1 Tysnd1 Cpne1 Zswim7 Ski Ccnd2 Tgfa Cst3 Fadd Hnrnpu Mdk Anapc2 Rnpep Nr3c1 Lrp5 Son Rgs2 Hnrnpk Asb2 Ctsb Larp4 Txn Smad7 Mapk1 Erbb4 Malat1 Tbl1x Kctd21 Itm2b Fli1 Eif3b Stt3b Hsp90b1 Ltbp1 Vps35 Rpl13a Ilf3 Ppp1r2 Aurkaip1 Apoe Ireb2 Rps11 Slc46a1 Mylip Abca1 Wiz Ctss Rpl35 Ncbp2 Rps8 Eef2 Kctd6 Rpl36 Hexb B2m Fbxw11 Ece1 Mtrf1 Rpl6 Cyfip1 Amz1 Ntan1 Dpp7 Rbm3 Eif3m
B
AAV−Syn−GFP
AAV−Syn−Cre
Egr1
Gap43
Cap23
Atf2
Mapkapk2
Vav2
Pcna
Rapgef4
Syt4
C
Klf4
Pten
Socs3
log2 Counts
Time after SCI (days)
0 1 3 7
*

**Figure S3. Differential gene expression analysis on RNA-seq of injured cortical neurons comparing wild-type and REST-depleted mice, related to Figure 3. (A)** Top gene ontology (GO) terms associated with DEGs (FDR p value < 0.1) comparing sensorimotor cortical neurons with or without REST depletion at day 7 following SCI. Bars represent FDR adjusted enrichment p-value in the negative log scale. Numbers in the bars indicate the number of DEGs overlapping existing in each GO term. The network plot shows DEGs enriched in each of the top up- or down-regulated GO terms. Colors indicates logFC of these genes comparing REST knock-out neurons to wild-type controls. **(B-C)** Expression levels of regeneration associated transcription factors (TFs) and genes comparing wild-type with REST knock-out cortical neurons recovered following SCI.

#### Figure S4

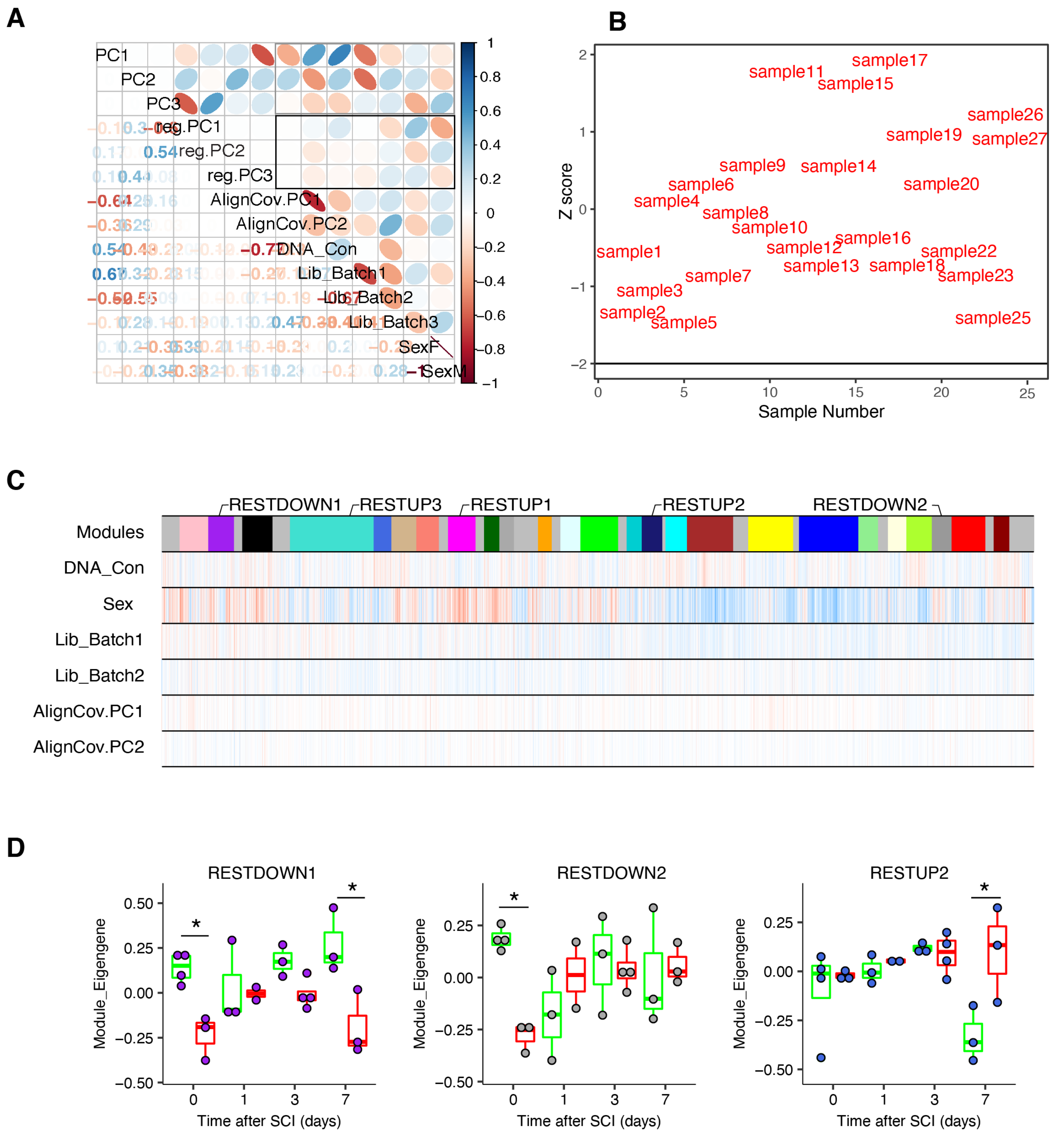
A
B
C
D
PC1
PC2
PC3
reg.PC1
reg.PC2
reg.PC3
AlignCov.PC1
AlignCov.PC2
DNA_Con
Lib_Batch1
Lib_Batch2
Lib_Batch3
SexF
SexM
Z score
Sample Number
Modules
Sex
RESTDOWN1
RESTUP3
RESTUP1
RESTUP2
RESTDOWN2
Module_Eigengene
Time after SCI (days)

**Figure S4. WGCNA analysis on RNA-seq of wild-type or REST knockout cortical motor neurons recovered following SCI, related to Figure 4. (A)** Correlations between the first three principle components (PCs) of gene expression data with sex, batch, library concentration and sequencing bias, before or after (highlighted by the black square) linear regression of these covariates from the RNA-seq data. **(B)** Sample connectivity to determine outliers and samples with |Z-score| < 2 were removed. **(C)** WGCNA module correlations with covariates. **(D)** Trajectory of the RESTUP1, RESTDOWN1 and RESTDOWN2 module eigengenes (MEs) across different time points after SCI in AAV-Syn-GFP (green) and AAV-Syn-CRE expressing (red) cortical neurons. Asterisks denote statistical significance assed by ANOVA model with Tuckey's post-hoc test *$p < 0.05$, **$p < 0.01$ comparing AAV-Syn-CRE to AAV-Syn-GFP. MEs and gene membership in each module (kME) are in Supplemental Table S3.

#### Figure S5

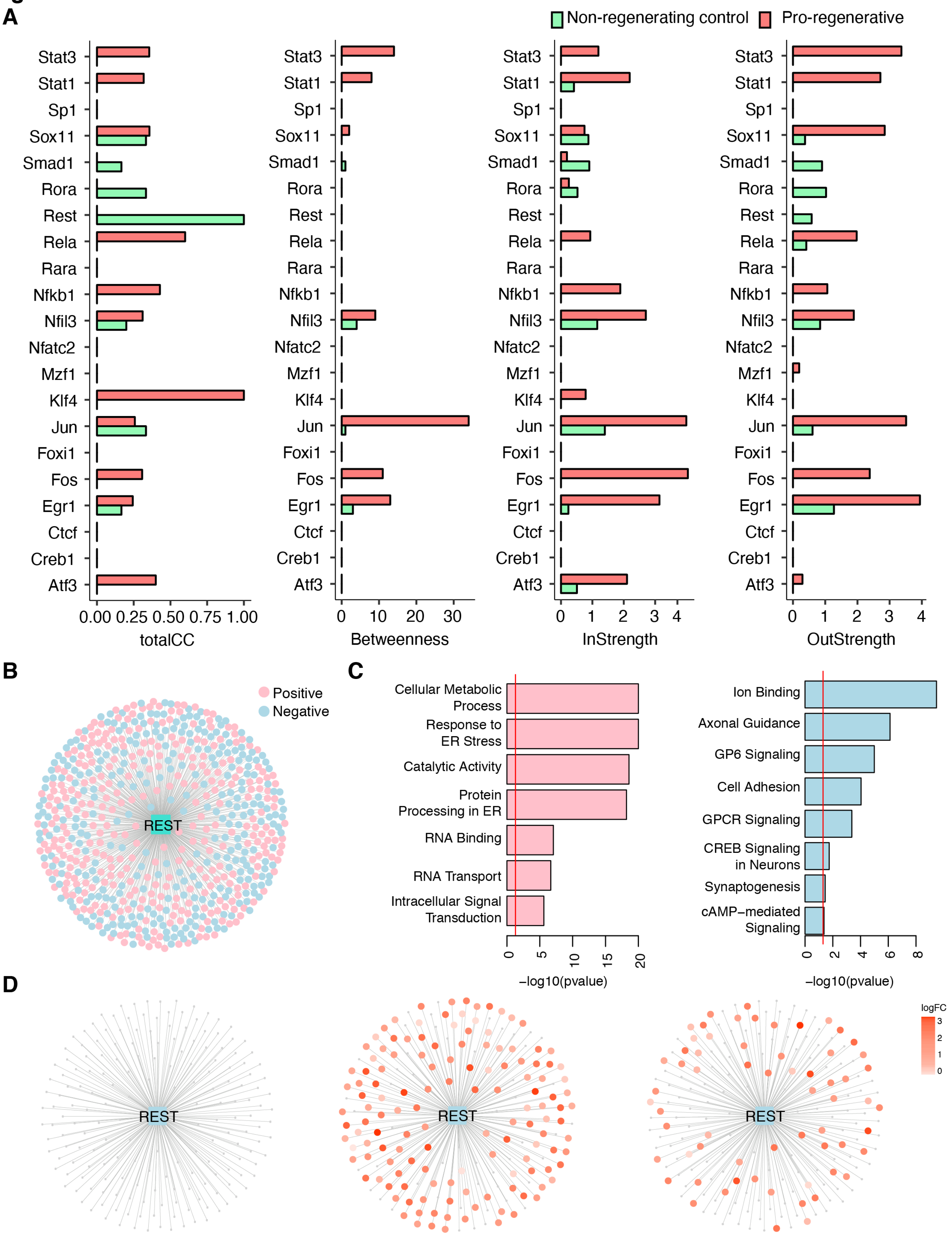

A
Non-regenerating control
Pro-regenerative
Stat3
Stat1
Sp1
Sox11
Smad1
Rora
Rest
Rela
Rara
Nfkb1
Nfil3
Nfatc2
Mzf1
Klf4
Jun
Foxi1
Fos
Egr1
Ctcf
Creb1
Atf3
0.00 0.25 0.50 0.75 1.00
totalCC
0 10 20 30
Betweenness
0 1 2 3 4
InStrength
0 1 2 3 4
OutStrength
B
Positive
Negative
REST
C
Cellular Metabolic Process
Response to ER Stress
Catalytic Activity
Protein Processing in ER
RNA Binding
RNA Transport
Intracellular Signal Transduction
0 5 10 15 20
−log10(pvalue)
Ion Binding
Axonal Guidance
GP6 Signaling
Cell Adhesion
GPCR Signaling
CREB Signaling in Neurons
Synaptogenesis
cAMP−mediated Signaling
0 2 4 6 8
−log10(pvalue)
D
REST
REST
REST
logFC
3
2
1
0

**Figure S5. TF networks comparing injured RGCs with pro-regenerative treatments or non-regenerative control, related to Figure 5.** TF networks were constructed from RNA-seq of RGCs sorted at 1, 3, 5 days after optic nerve crush alone (non-regenerating control) or with pro-regenerative treatment (AAV2-shPten.mCherry/Oncomodulin/CPT-cAMP; pro-regenerative), using the unbiased, step-wise pipeline described in Figure 2A. **(A)** Parameters indicating connectivity of each TF in the networks of pro-regenerative RGCs and non-regenerating control RGCs. **Local clustering coefficient (totalCC):** the clustering coefficient of a vertex which is calculated as the number of links between the vertices within its neighborhood divided by the number of links that are possible between them. A high local clustering coefficient suggest a tightly connected local network. **Betweenness**: the fraction of the shortest paths between all pairs of vertices that pass through one vertex. **InStrength or OutStrength:** the number of links connected to one vertex. In directed and weighted networks, the number of arcs that end at the node is defined as "InStrenght", and the number of arcs that start from the node is defined as "OutStrength". **(B)** Genes that are positively regulated (activated, pink-colored nodes) or negatively regulated (repressed, blue-colored nodes) by REST defined by ARACNe from RNA-seq of RGCs sorted at 1, 3, 5 days after optic nerve crush alone or with pro-regenerative treatment. FDR adjusted p value $< 0.05$, permutations = 100, bootstrap consensus = 95% were used to identify REST-regulated genes by ARACNe. **(C)** Top GO terms of REST-activated genes and REST-repressed genes. **(D)** Overlap between REST-repressed genes and up-regulated genes by pro-regenerative treatment in RGCs at day 3 and day 5 following optic nerve crush. Colors represent logFC of significantly up-regulated genes by pro-regenerative treatments determined by FDR $< 0.1$. A full list of REST-repressed genes is shown in Supplemental Table S4.

### Figure S6

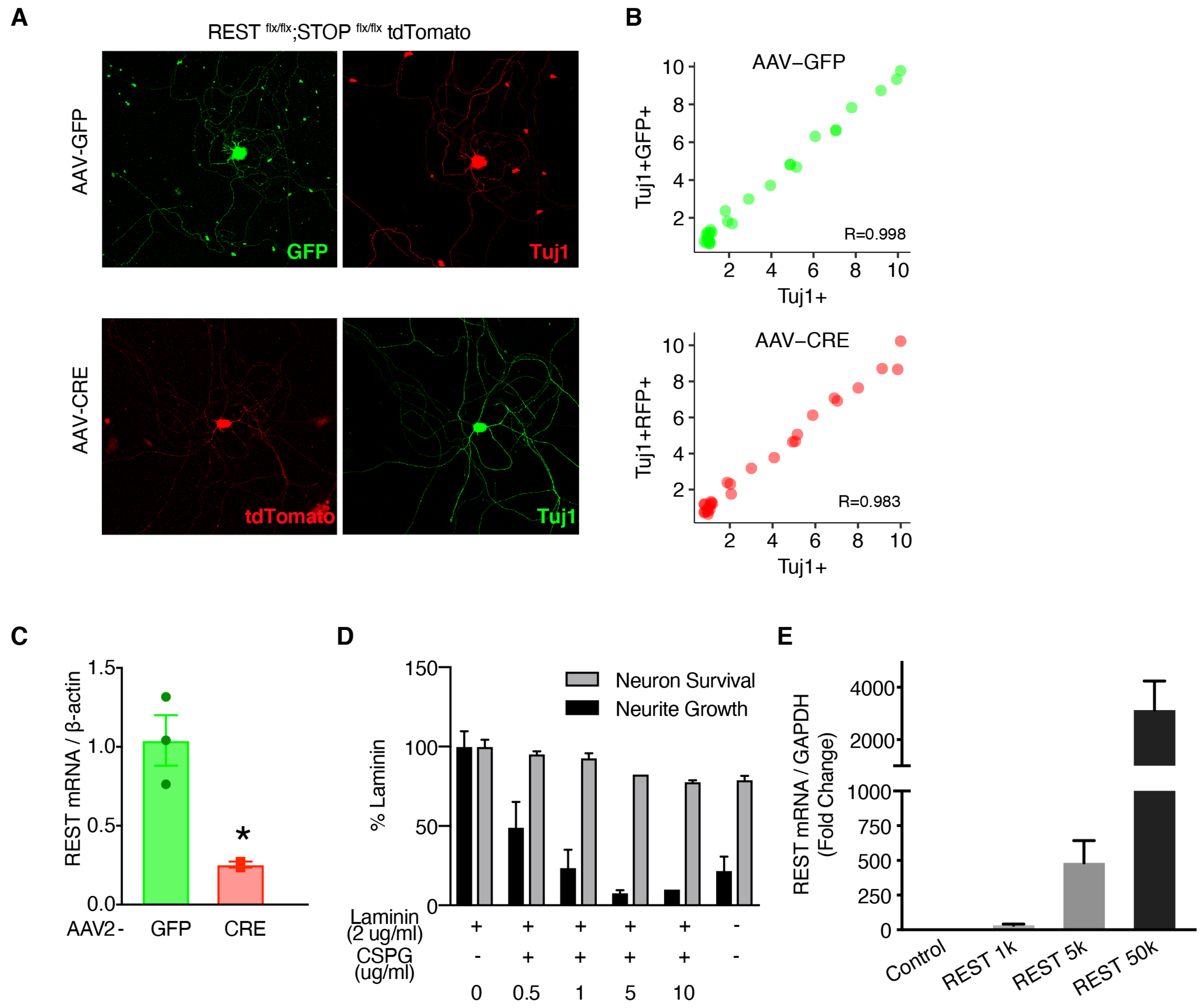
A
REST flx/flx;STOP flx/flx tdTomato
AAV-GFP
GFP
Tuj1
AAV-CRE
tdTomato
Tuj1
B
AAV−GFP
Tuj1+GFP+
Tuj1+
R=0.998
AAV−CRE
Tuj1+RFP+
Tuj1+
R=0.983
C
REST mRNA / β-actin
AAV2 -
GFP
CRE
*
D
% Laminin
Neuron Survival
Neurite Growth
Laminin
(2 ug/ml)
CSPG
(ug/ml)
0
0.5
1
5
10
E
REST mRNA / GAPDH
(Fold Change)
Control
REST 1k
REST 5k
REST 50k

**Figure S6. Validation of REST overexpression or knockdown in vitro, related to Figure 6.** REST flx/flx;tdTomato DRG cells were transduced with AAV-GFP or AAV-CRE for 7 days. **(A)** Confocal images of GFP or tdTomato co-stained with the neuronal marker Tuj1 to confirm efficient AAV transduction into DRG neurons as well as Cre-induced tdTomato expression, an indication of REST deletion. **(B)** A Pearson correlation between the number of Tuj-expressing neurons (Tuj1+) and the ones also expressing GFP (Tuj1+GFP+; AAV-GFP transduced neurons) or tdTomato (Tuj1+RFP+; AAV-CRE transduced neurons). Each dot represents an individual image quantified. A high correlation suggests efficient transduction of AAVs into the DRG neurons. **(C)** qRT-PCR of REST mRNA levels to confirm REST knockdown. **(D)** Neurite outgrowth and neuronal survival at indicated doses of CSPG to determine an optimal dose of CSPG used in Figure 6A. **(E)** qRT-PCR of REST mRNA levels in DRG neurons to confirm REST overexpression by transducing the lentiviral constructs containing REST. Humanized luciferase protein (Lv135-hLuc) was used as a control. Bars represents means±SEM for **(C-E)**.

#### Figure S7

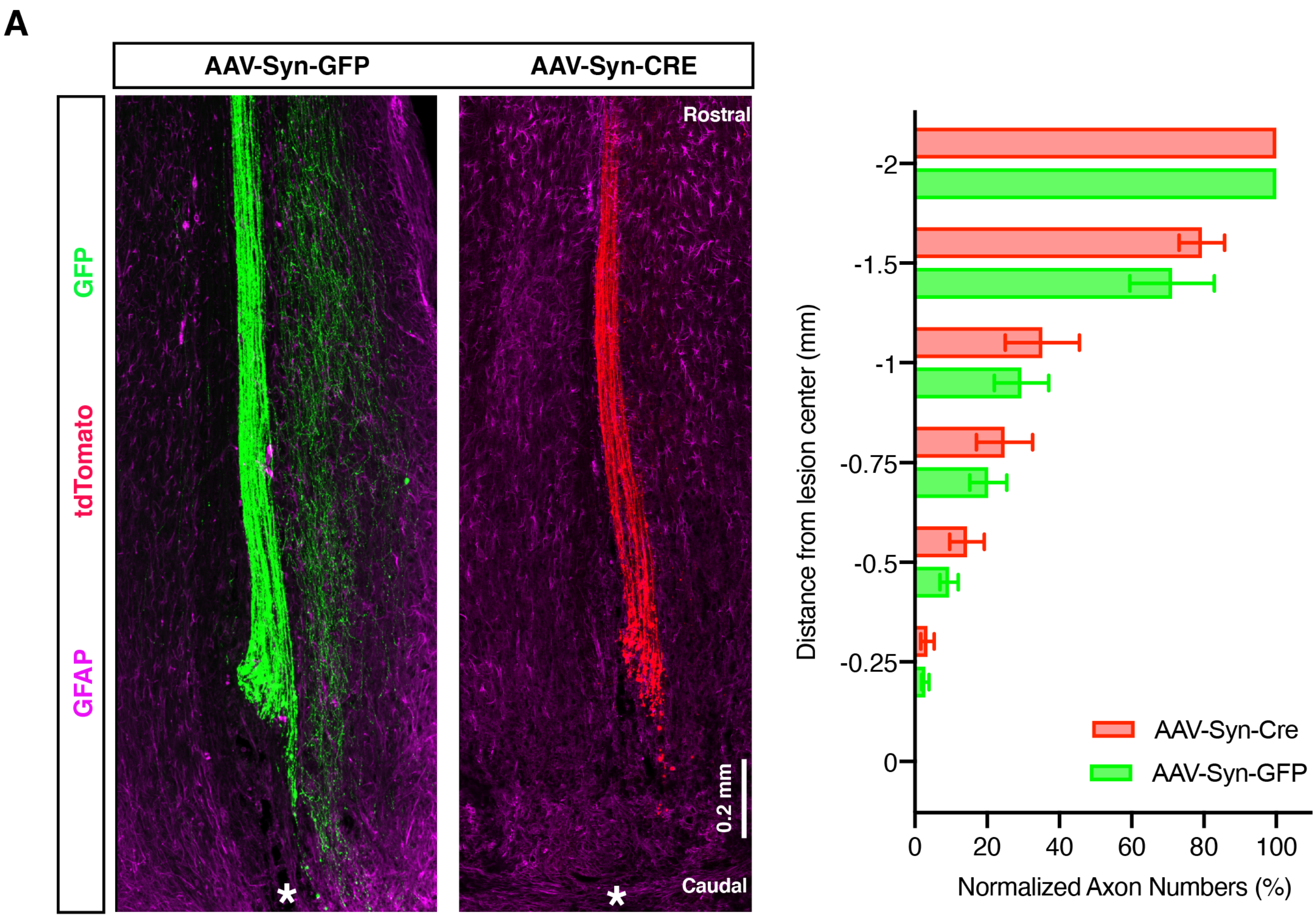
A
AAV-Syn-GFP
AAV-Syn-CRE
GFP
tdTomato
GFAP
Rostral
Caudal
0.2 mm
*
-2
-1.5
-1
-0.75
-0.5
-0.25
0
Distance from lesion center (mm)
0
20
40
60
80
100
Normalized Axon Numbers (%)
AAV-Syn-Cre
AAV-Syn-GFP

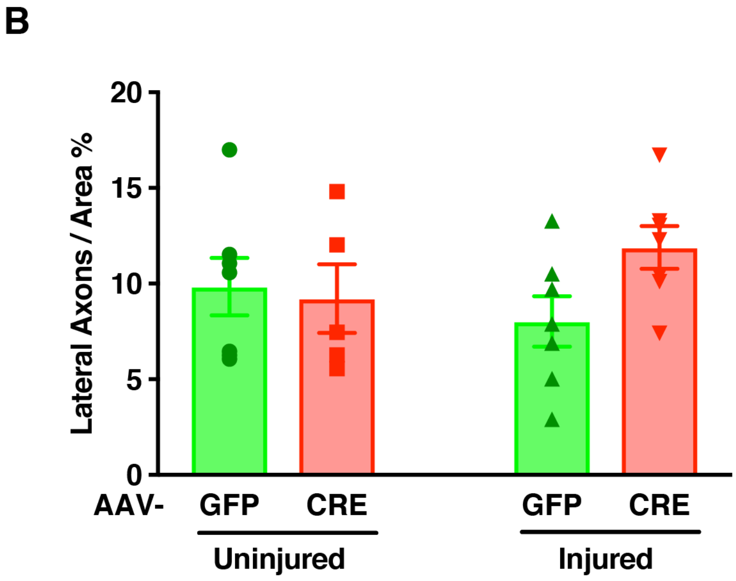
B
20
15
10
5
0
Lateral Axons / Area %
AAV-
GFP
CRE
GFP
CRE
Uninjured
Injured

**Figure S7. Corticospinal (CST) axon staining 3 days after spinal cord injury or in non-injured conditions, related to Figure 7. (A)** Confocal images and quantitation of GFP or tdTomato labeled axons in horizontal sections of lesioned spinal cord also stained for astrocytes (glial fibrillary acidic protein (GFAP)). REST $^{flx/flx}$;tdTomato mice were injected into the sensorimotor cortex with AAV-Syn-GFP (wild-type) and AAV-Syn-CRE (REST cKO) for 4 weeks followed by a full crush at thoracic spinal cord level 10 (T10). Spinal cord was recovered 3 days post-injury. Lesion center was marked with *. To quantify axon numbers at indicated distances from lesion center, N=4 mice for each group was used. Bars represent means±SEM; no statistical difference was found between AAV-Syn-GFP with AAV-Syn-Cre using two-way ANOVA followed by Sidak. **(B)** Quantitation of total number of BDA-labeled axons in uninjured, or injured REST $^{flx/flx}$ mice receiving AAV-GFP or AAV-CRE. The paradigm for AAV transduction, spinal cord injury and BDA labeling was the same as described in Figure 7, except that SCI was not performed on uninjured mice. Total number of BDA-labeled axons in injured mice was counted 3 mm rostral to the lesion center. Bars represent means±SEM; No statistical difference was found by one-way ANOVA with Sidak post-hoc test compared to uninjured AAV-GFP.

### Figure S8

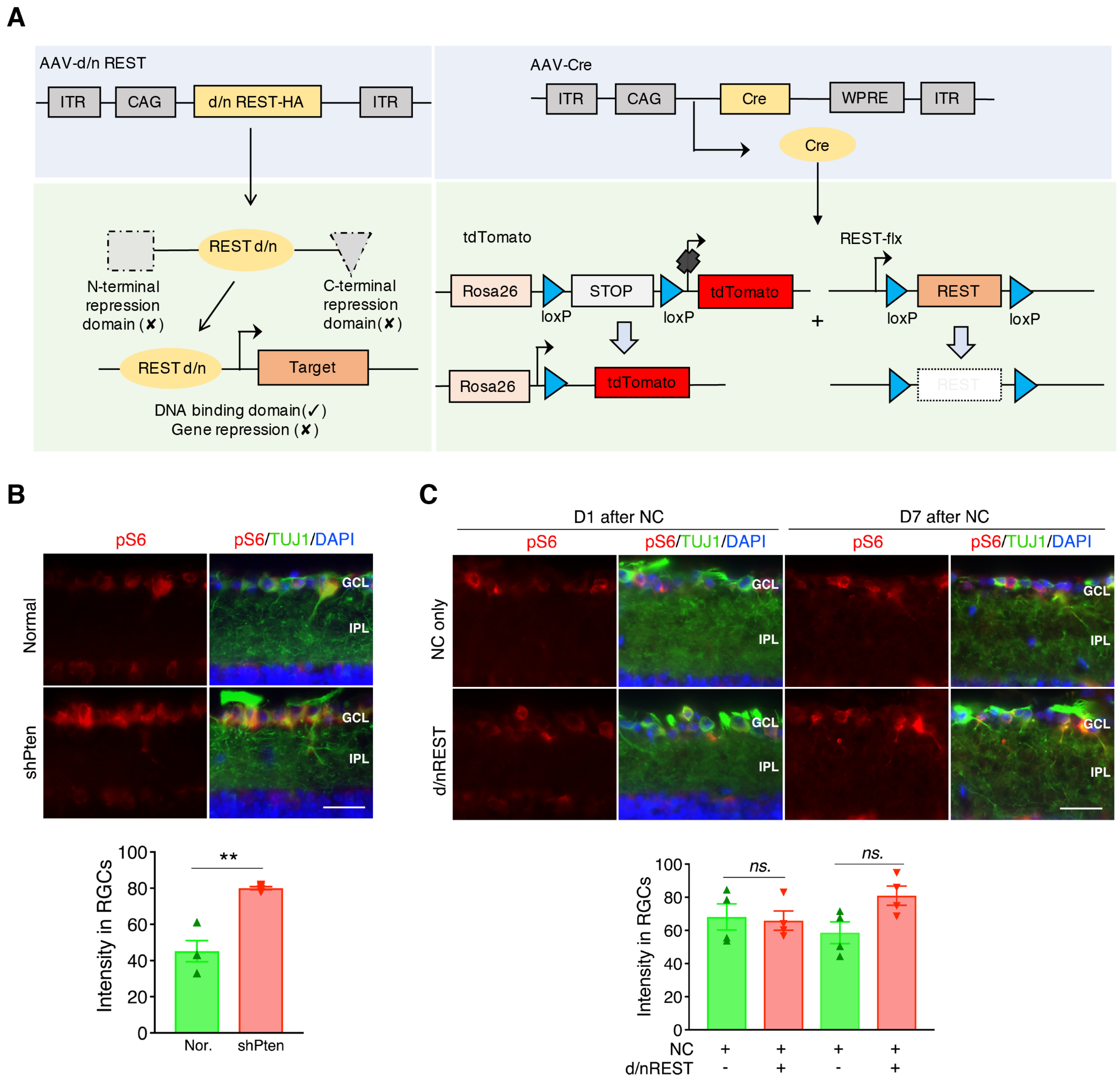
A
AAV-d/n REST
ITR
CAG
d/n REST-HA
ITR
AAV-Cre
ITR
CAG
Cre
WPRE
ITR
Cre
REST d/n
N-terminal repression domain (✘)
C-terminal repression domain (✘)
REST d/n
Target
DNA binding domain (✓)
Gene repression (✘)
tdTomato
Rosa26
STOP
tdTomato
loxP
loxP
Rosa26
tdTomato
REST-flx
REST
loxP
loxP
+
B
pS6
pS6/TUJ1/DAPI
Normal
shPten
GCL
IPL
Intensity in RGCs
**
Nor.
shPten
C
D1 after NC
D7 after NC
pS6
pS6/TUJ1/DAPI
NC only
d/nREST
GCL
IPL
Intensity in RGCs
ns.
NC
d/nREST

**Figure S8. Conditional depletion or inhibition of REST in the model of optic nerve injury, related to Figure 8. (A)** Schematic displaying strategies to induce REST inhibition or REST depletion in retina cells. **(B-C**) Inactivation of PTEN or REST initiates different downstream signaling pathways. Changes of phospho-S6 immunostaining (pS6, red) in RGCs (visualized by immunostaining for b**III** tubulin, green) after AAV2- mediated knock-down of PTEN (shPten) after optic nerve crush (NC), comparing with normal retina (Nor.) **(B)** or expression of dominant-negative REST (d/nREST) **(C)**. Whereas knockdown of PTEN elevated levels of p-S6 **(B)**, REST deletion did not (**C**: expression shown at 1 and 7 days after nerve crush). ** $p < 0.01$ *t*-test in A, one-way ANOVA followed by Sidak test in B. Scale bars: 45 µm.
